# Population Structure and Migration in the Eastern Highlands of Papua New Guinea; a Region Impacted by the kuru Epidemic

**DOI:** 10.1101/2022.09.26.509478

**Authors:** Liam Quinn, Ida Moltke, Jerome Whitfield, Michael Alpers, Tracy Campbell, Holger Hummerich, William Pomat, Peter Siba, George Koki, John Collinge, Garrett Hellenthal, Simon Mead

## Abstract

Populations of the Eastern Highlands of Papua New Guinea (EHPNG, area 11,157 km^2^) lived in relative isolation from the rest of the world until the mid-20^th^ century, and the region contains a wealth of linguistic and cultural diversity. Notably, several populations of EHPNG were devastated by an epidemic prion disease, kuru, which at its peak in the mid-twentieth century led to some villages being almost depleted of adult women. Until now, population genetic analyses to learn about genetic diversity, migration, admixture and the impact of the kuru epidemic have been restricted to a small number of variants or samples. Here, we present a population genetic analysis of the region based on genome-wide genotype data of 943 individuals from 21 linguistic groups and 68 villages in EHPNG, including 34 villages in the South Fore linguistic group, which the group most affected by kuru. We find a striking degree of genetic population structure in the relatively small region (average F_ST_ between linguistic groups 0.024; area similar to Jamaica). The genetic population structure correlates well with linguistic grouping, with some noticeable exceptions. Also, we find evidence of a series of discrete admixture events that appear to coincide with previously expected introduction dates and movement of the sweet potato in the region. Finally, we find signatures of more recent migration within the EHPNG region and observe a substantial excess of female migration into the heavily kuru-affected South Fore linguistic group (p=0.0017, Chi Squared Test), likely reflecting the sex-bias in incidence of kuru. These data provide an in-depth look at the population genetics of a region devastated by a prion disease epidemic and which was until recently relatively isolated from the rest of the world.

## Introduction

The Eastern Highland region of Papua New Guinea (EHPNG) that covers an area of 11,157 km^2^ was more or less isolated from the rest of the world until the early decades of the 20^th^ century(1). At the beginning of the 20^th^ century, western observers held the belief that the highlands of New Guinea were likely uninhabited(2, 3). Exploration by Christian missionaries and gold prospectors, however, revealed the highlands to contain heavily populated valleys that were home to close to one million people(2, 4). From 1918 until its independence in 1975 the region was gradually subjected to Australian colonial rule(5). Colonial authorities divided the peoples of EHPNG into administrative groups based on the language spoken in particular areas and to date, most studies of the people in EHPNG and the PNG Highlands in general have used these linguistic groupings as convenient population labels(1).

The people of EHPNG are notable for their complex cultural(6) and trade systems(7), cosmology, and linguistic diversity(1, 8). The number of separate languages spoken in the broader highlands of PNG is believed to be in the hundreds and in the EHPNG region alone there are 37 distinct linguistic groups(1). Historically, each linguistic groups consisted of clans(9, 10), with clan composition dynamic, and changed depending on clan disputes, wars and other cultural factors(11). The region has extreme terrain including mountains, valleys and fast-flowing river systems that impeded travel(1, 7).Yet, anthropological studies have pointed to a complex picture of migration of individuals and groups over both long and short distances within EHPNG(11, 12). In addition to this, theories abound about possible admixture within EHPNG from other fields of study: linguists, archaeologists and historians have discussed competing theories as to how the region was originally populated and the origin of the current groupings(13-15). The presence of non-indigenous products in EHPNG evidence long distance trade networks, proving EHPNG not to be completely isolated, and that trade may have led to genetic exchange(1). Furthermore, there are hypotheses of migration both within and into the region in the literature (16, 17). Genetic data could help reveal how factors like linguistic diversity, cultural and agricultural practices, migration and extreme geography can shape population genetic structure in a region that was until recently relatively isolated from the rest of the world.

EHPNG is also known because several of the linguistic groups in the region (mainly the Fore and groups with whom they intermarried) were devastated by an epidemic of the prion disease kuru during the 20^th^ century(18, 19). Prion diseases are fatal transmissible neurodegenerative disorders caused by the propagation of prions, infectious agents composed of multichain assemblies of misfolded prion protein(20). Kuru was transmitted by endocannibalistic mortuary feasts at which a deceased relative was consumed by kith and kin in a ritualistic manner(6, 9). The epidemic resulted in close to 2,700 recorded deaths and predominantly affected adult women and children of both sexes because they consumed the most highly prion-infected tissues(19). During the height of the epidemic observers noted many villages were depleted of adult women(21). Hence the epidemic likely had an impact on established cultural and demographic processes, like migration in the region(5). Moreover, kuru imposed strong selection pressure on the affected populations, with evidence of strong balancing selection acting at the prion protein gene locus(21). Whether kuru imposed selection pressure at other loci is currently unknown, but a prerequisite of answering this question is a deeper understanding of the population genetics of the region.

To date relatively few genetic studies of the EHPNG region have been performed(17, 22, 23). These have shown that groups that populate the broader highlands region of PNG display high levels of genetic differentiation(16, 17). However, no studies have yet had enough data from EHPNG to fully explore population diversity, structure and migration in this subregion of the highlands, and its correlations with language, topography and kuru disease. Motivated by this, we analysed genetic data for 943 individuals from EHPNG sampled from 21 of the 37 distinct linguistic groups in the region. Moreover, with dense sampling from a total of 68 villages we have fine-scale village level data for several of the linguistic groups. We present an in-depth analysis of the population structure and admixture in EHPNG that reveals a striking complexity of the small region. Subsequently we apply knowledge of this population structure to address questions regarding the population genetic impact of kuru. Finally, we discuss the likely causes of this complex population structure, what it can tell us about the history and population dynamics of the region and what this might entail for potential future studies of EHPNG and of kuru.

## Materials and methods

### Ethical aspects

Laboratory studies were approved by the Papua New Guinea Medical Research Advisory Committee, and by the local research ethics committee of UCL Institute of Neurology. Participation of the communities involved was established and maintained through discussions with village leaders, communities, families and individuals. The field studies followed the principles and practice of the Papua New Guinea Institute of Medical Research (PNGIMR) which included individual oral consent from all participants before any samples were obtained.

### EHPNG samples and genome-wide genotyping

Blood samples were taken from 4,217 individuals from communities in EHPNG by members of PNGIMR. Information was obtained about the individual’s village of residence, and sex. After initial processing locally these samples were transported to the Medical Research Council Prion Unit (MRCPU) in the United Kingdom. The samples were then further processed and genotyped in several stages. The first 488 samples were genotyped on the Illumina 670 genotyping platform. The following 1,106 samples were genotyped on the Illumina Omni Express genotyping platform. 83 individuals were genotyped on both the Illumina 670 (678,000 variants) and Illumina Omni Express (748,000 variants) platforms to permit downstream analysis of possible biases introduced by the different genotyping platforms.

### Other samples and genotype data used for context

In addition to the EHPNG samples we also included samples from previous studies in a subset of the analyses. Specifically, we included 380 samples(17) which consisted of samples from 84 PNG linguistic groups and surrounding islands. Among these samples were 26 individuals from 10 EHPNG linguistic groups. All of these 380 samples were genotyped using the Ilumina Infinium Multi-Ethnic Global Array. We also included samples from the CEU (Americans of European descent), CHB (Han Chinese) and YRI (Yoruba Nigerian) populations from the 1000 genomes project. For these populations we downloaded phase 3 Omni genotyped data from the International Genome Sample Resource portal (https://www.internationalgenome.org/category/omni/)(24).

### Other samples and genotype data used for phasing

The primary dataset was incorporated with other available data to create a phasing panel with as much population breadth as possible. In addition to the EHPNG data, 1000 genomes phase 3 data, data obtained through separate group access agreement including 34 African populations, 2 indigenous South American populations, 4 ancient European individuals, a Denisovan and a Neanderthal individual were merged. This gave a haplotype phasing panel of 4,779 individuals (See Supplementary Table 1c for breakdown).

### Quality filtering and phasing of the EHPNG data

After merging the EHPNG data with the phasing panel dataset, quality control was performed in PLINK 1.9(25). Specifically, we first filtered SNPs with allele frequency below 1% and individuals with missingness above 2%. This resulted in 4,479 individuals, including 1,374 from EHPNG and 122,662 variants available for phasing. We then jointly phased the autosomal chromosomes for all individuals using SHAPEIT(26) with default parameters and the linkage disequilibrium-based genetic map build 37.

### Creation of datasets for genetic analysis and relatedness checks

Three datasets were created for subsequent analysis through subsetting individuals from the phased data. The first dataset was designed to analyse population structure in EHPNG based on linguistic group membership and without possible impacts of sampling size bias or genotyping platform and it comprised of 16 individuals from 20 linguistic groups to give 320 individuals, who were all originally genotyped on the Illumina Omni Express platform. This dataset is referred to as the *Linguistic Group* dataset. Before randomly selecting 16 individuals from each of the 20 linguistic groups a relatedness threshold was applied in plink 1.9 of PIHAT < 0.2 (using the -genome option after data for each linguistic group was linkage disequilibrium pruned with the command –indep-pair-wise 50 5 0.4 to extract between 70,000-75,000 independent markers for relatedness analysis).

The second dataset was compiled to examine effects at the inter-village level using all the EHPNG samples, referred to subsequently as the *Village Analysis* dataset. For this analysis a relatedness threshold of 0.1875 PIHAT was applied to each of the linguistic groupings with linkage disequilibrium pruning done identically for each linguistic group as per the *Linguistic Group* dataset. This gave 943 EHPNG individuals from 21 linguistic groups and from 68 different villages (see Supplementary Table 1a for breakdown).

The third dataset was used with the intention of examining the relationship of EHPNG individuals and population clusters defined in the *Village Analysis* dataset analysis in the context of other PNG samples. To create this dataset, we merged the *Village Analysis* dataset with the data from 380 individuals from PNG and surrounding islands. This is referred to subsequently as the *External Analysis* dataset.

In addition to the three datasets described above we also created a number of datasets for ADMIXTURE analyses with the purpose of investigating possible admixture due to recent contact between EHPNG individuals and individuals with European ancestry. To create these, we merged unphased EHPNG data (See Supplementary Table 6 for breakdown) from each linguistic group separately with individuals from the 1000 genomes populations CEU (Americans with European ancestry, n=100) and CHB (Han Chinese individuals, n=105).

### Principal Component Analysis (PCA), Fixation Index (F_ST_), heterozygosity runs of homozygosity (ROH), linkage disequilibrium (LD) and relatedness estimation of *Linguistic group* dataset individuals

We performed PCA using the PLINK 1.9 command “—pca”, on the *Linguistic group* dataset. PCA analysis highlighted 7 individuals who clustered distantly from others in their own linguistic groups and were removed from subsequent F_ST_, LD, ROH, heterozygosity and relatedness calculations to avoid any potential effect of potential recent ‘long distance’ migration in the region.

Pairwise F_ST_ calculations were performed between linguistic groups of the remaining 313 individuals from the 20 linguistic groups. This was done in plink 1.9 using the --fst command after linkage disequilibrium pruning of the data following the same procedure as for PCA.

For calculation of heterozygosity, relatedness, ROH and LD all linguistic groups were downsampled to 13 individuals to give equal numbers of individuals per linguistic group as these statistics are impacted by sampling size differences. For heterozygosity estimations, given the heterogeneity observed between linguistic groups in PCA and F_ST_ analyses we decided to include all SNPs available in the Illumina Omni merge to avoid any ascertainment bias issues. Heterozygosity was measured by performing the –freq command and then calculating the genome wide average of 2pq per locus where p is the major allele frequency and q is the minor allele frequency. ROH were estimated for each of the linguistic groups in the analysis. This was done using PLINK 1.9 and the –homozyg setting. A sliding window of 50 SNPs was used and a SNP density of 50 SNPs per kilobase. Counts of ROH were based on any ROH being a minimum of 10mb in length.

LD decay curves were created for each of the linguistic groups. This was performed using data from chromosome 22 and the commands (--ld-window 1000000, --ld-window-kb 600, --ld-window-r2 0, -- maf 0.05, --r2 dprime with-freqs). The same analyses were also performed on 13 randomly chosen individuals from each of the 1000 Genomes populations CEU and YRI (Yoruba Nigeria) to allow comparison to other population outside PNG.

Overall relatedness calculations were performed to check to what extent recent close relatedness of individuals was impacting this comparative analysis of summary genetic data. Average PIHAT between individuals was calculated for each of the 20 linguistic groups. This was again using linkage disequilibrium pruned data (the same procedure as done for relatedness estimation to when creating the datasets).

### ChromoPainter and fineStructure Analyses

We applied ChromoPainter(27) (CP) on each of the *Linguistics Group, Village Analysis*, and *External Analysis* datasets to identify patterns of shared ancestry between individuals in the datasets. CP is a ‘painting’ technique that compares haplotype patterns within a target chromosome to those within a set of sampled “donor” chromosomes. In a genetic region, if a target’s haplotype patterns are more similar to a particular donor relative to the other donors, this suggests the target shares a more recent ancestor with that donor relative to the others for that genetic region. CP provides a ‘painting profile’ for each target individual reflecting the amount of genome-wide DNA for which the individual is inferred to share a most recent ancestor with each donor individual.

Prior to each of the analyses, we first estimated two CP model parameters, the switch (“-n”) and emission (“-M”) rates, using 10 Expectation-Maximisation (E-M) iterations (“-i 10 -in -iM”) for chromosomes 1, 5, 7, 15, and 21. We then fixed these estimated values and separately painted each of the individuals in each dataset as a recipient when using the other individuals in the same dataset as donors.

We next used fineStructure(27) (FS) to group individuals into genetically homogenous clusters based on CP output of the three datasets. Importantly, these groupings are free from any bias due to a priori classifications of individuals, e.g. based on linguistic group classifications. Following the recommended FS approach described by a previous publication(27), we inferred a normalisation parameter ‘c’ and performed two million iterations of Markov-Chain-Monte-Carlo (MCMC), sampling an inferred clustering every 10,000 iterations after a burn in of one million iterations. Starting from the single MCMC sampled clustering with highest posterior probability, we then performed 100,000 additional hill-climbing steps in FS to find a nearby state with even higher posterior probability. The results of this hill-climbing approach grouped individuals into homogenous clusters based on the highest maximum likelihood. FS creates a bifurcating tree that presents the relationship of these clusters to one another. A next-neighbour joining algorithm was used to ascend the FS-derived tree to obtain a more interpretable number of clusters that could best describe overall population structure in the region.

External non-EHPNG individuals were grouped together based on FS clustering of the *External Analysis* CP copy vector matrix. Clusters that were difficult to interpret and very heterogeneous were eliminated from SF analysis. This resulted in 13 external population clusters being used in the analysis (Supplementary Table 4a for composition of 13 external population clusters).

### ADMIXTURE

We tested for admixture between EHPNG linguistic groups and 1000 genomes CEU and CHB populations using ADMIXTURE(28) with K=2. The analyses of each dataset were run 5 times to ensure convergence was reached.

### SOURCEFIND

We used SOURCEFIND to form the painting profile of each of the 13 *EHPNG Clusters* as a mixture of those of other groups (29). SOURCEFIND uses a Bayesian approach that puts a prior on the number of groups contributing >0% to this mixture, hence eliminating contributions that cannot be reliably distinguished from background noise. For this analysis, we used the painting profiles from the *External Analysis* CP painting (Supplementary Figure 6).

We used SF to form each EHPNG cluster as a mixture of (i) 13 non-EHPNG clusters (defined in FS section) or (ii) 13 non-EHPNG clusters and the 12 other EHPNG clusters. The first analysis explores the extent to which each EHPNG shares recent ancestry with groups outside the EHPNG, while the second explores recent ancestry sharing among EHPNG groups.

### GLOBETROTTER

We applied GLOBETROTTER(30) (GT) to infer admixture events in each of the 13 *EHPNG Cluster* populations under a “pulse” model whereby admixture occurs instantaneously for each admixture event, followed by the random mating of individuals within the admixed population from the time of admixture until present-day. To do so, we used the CP External analysis painting and used all XX non-EHPNG clusters as potential surrogates to the admixing sources, letting GT identify which ones are the best representatives of these unknown sources.

### Testing for sex-biased migration in South Fore linguistic group

To test if there is a sex-bias among the migrants in the highly kuru affected South Fore linguistic group we identified migrants among 331 South Fore individuals. Migrants were classified using FS clustering of the *Village Analysis dataset*. Classification of migrants was influenced by our understanding of population structure at the regional scale (Figure 1) and at a fine-scale (Figure 2). Specifically, migrants into the South Fore linguistic group were defined as any South Fore individuals who were classified into one of the 11 EPHNG clusters besides *Clusters 5* and *7* (Supplementary Table 3) that contain the vast majority (82.9%) of South Fore individuals. However, we made two exceptions based on the observation in our fine-scale structure analyses that some villages situated close to linguistic group borders have FS clustering profiles that more closely resemble the group on the other side of the border: individuals were not classified as migrants if 1) they resided in the villages Ilesa, Awarosa and Amora, but were placed in *EHPNG Cluster 8* (Awa dominated) or 2) they were from villages in the South Fore bordering Gimi and North Fore, who were placed in *EHPNG Clusters 12* (Gimi) and *6* (North Fore). Among the 376 individuals from non-kuru affected linguistic groups we classified individuals as migrants if they were placed in clusters which comprised individuals from different linguistic groups to their own. We then used a chi-squared statistic to test whether there were more female migrants than expected given overall gender ratios in the South Fore. For comparison, we performed the same test in 15 linguistic groups (376 individuals) without a history of kuru.

**Figure 1.**
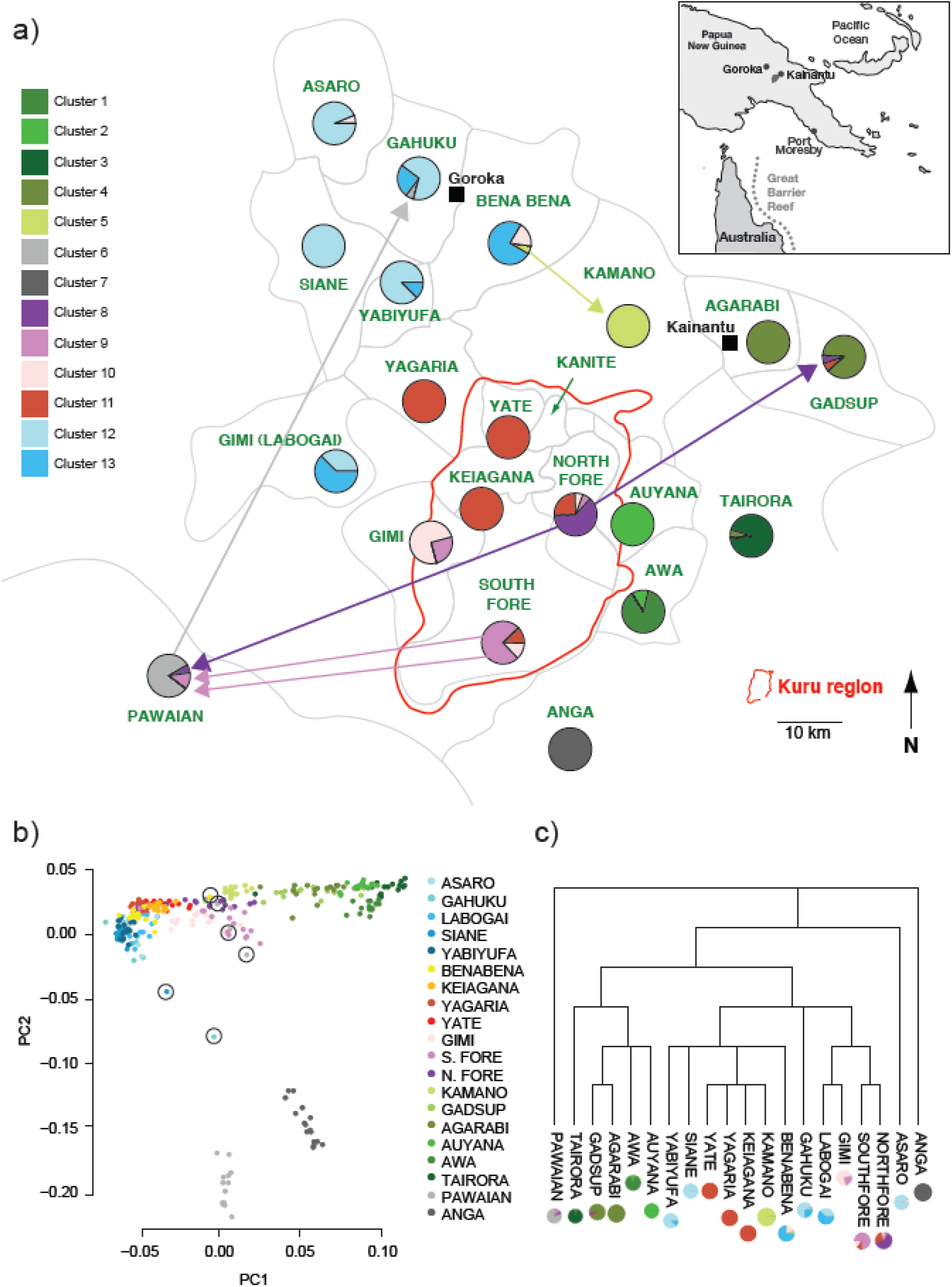
Population Structure of EHPNG described through linguistic group membership. 1a) Map of EHPNG region overlaid with results of FS clustering for 20 EHPNG linguistic groups. Arrows indicate individuals highlighted in PCA analysis who also cluster aberrantly from other individuals in the same linguistic group. b) PCA analysis of the same 320 individuals. Circled individuals are placed distantly from others from the same linguistic group. c) Best estimate language tree for 20 EHPNG linguistic groups (source http://glottolog.org) FS clustering profiles for each linguistic group are placed alongside their labels.

**Figure 2.**
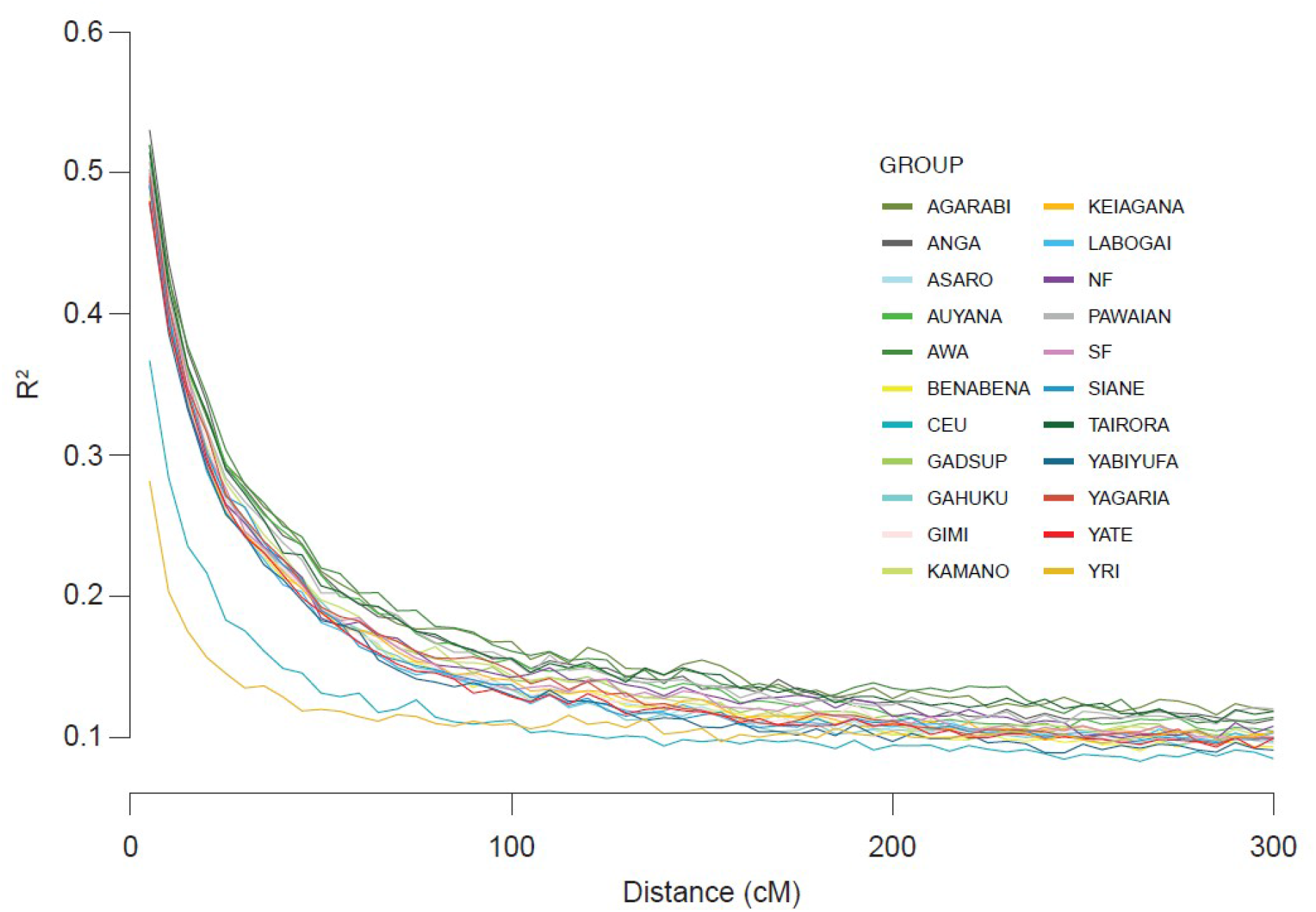
LD decay curves for 20 EHPNG linguistic groups. 13 individuals analysed per group, and 13 individuals from each of the 1000 genomes YRI and CEU populations whose curves are below the EHPNG curves.

## Results

### Analyses of Population Structure of EHPNG Region

To analyse population structure we used data from 20 linguistic groupings in EHPNG(1) (Figure 1a). To control for effects of sampling size(31) we carried out all of these analyses out on a reduced dataset of 320 individuals with 16 not closely related individuals from each of 20 linguistic groups, the *Linguistic Group Dataset* (see Materials and Methods).

First, we performed principal components analysis (PCA) and plotted the results with individuals coloured according to their linguistic group membership (Figure 1b). Principle Component (PC) 1 and PC2 shows individuals from the same linguistic group broadly cluster together. PC1 shows placement of linguistic groups along a North West – South East cline, whereas PC2, separates the Anga and Pawaian linguistic groups from the other groups. Notably, the linguistic groups that overlap on the plot tend to be geographically neighbouring linguistic groups, for example individuals from linguistic groups in the North-West of the region (Asaro, Siane, Yabiyufa, Labogai and Gahuku) cluster closely together. Examination of further PCs shows that several of the other linguistic groups separate out on some of the higher PC axes, including Tairora, Awa, Auyana, Kamano, Agarabi and Gadsup, all groups living in the Eastern part of the region (See Supplementary Figure 1 for PC3-10). Other linguistic groups, including the groups in the North-West do not entirely separate out within the first 10 PCs. It is also worth noting, that in the PC1-PC2 plot there are several outlier individuals including three Pawaian, one Gadsup, one Gahuku, one Siane, and one Kamano individual, who do not cluster with other individuals from the same linguistic group and may represent migrant individuals or descendants of migrants within the region.

We estimated genetic differences between groups using pairwise F_ST_ values from the *Linguistic Group dataset* after removing the clear outlier individuals observed on visual inspection of the PC1-PC2 plot (Supplementary Figure 2). On average, the pairs of linguistic groups have greater genetic differences than those found between groups of individuals from nations in Europe as distant as Finland and Spain (average F_ST_ in EHPNG 0.024 vs 0.012 for Finland and Spain (8)) despite being on average only 45km apart. Consistent with the PCA analysis results, the Anga and Pawaian groups had the largest average pairwise F_ST_ values to other groups (Anga average 0.046, range 0.039-0.060; Pawaian average 0.050, range 0.043-0.064). At the other end of the spectrum, smaller than average values were seen between the linguistic groups in the North-West of the region that grouped together in the PC1-PC2 plot (average 0.0037, range 0.0021-0.0057). Additionally, we observed very small values for a few of the geographically neighbouring group pairs like the South Fore and Gimi. Interestingly, the differences in F_ST_ seem to roughly follow a pattern of increasing group heterogeneity when moving through the region from North-West to South East (Supplementary Figure 2). More broadly, the affinities and differences between groups according to F_ST_, combined with the PCA results, seem to suggest genetic structure that roughly follows linguistic grouping but as importantly, the geographical sub-regions in EHPNG: North-West (blue colours in PCA), Upper-Mid (red colours in PCA), Lower-Mid (purple colours in PCA), North-East (light green colors in PCA), South-East (dark green colours in PCA) and South (gray colours in PCA), Pawaian and Anga. These geographical groupings of linguistic groups also represent linguistic groups that reside closely with one another on suggested linguistic trees of the region (Figure 1c for linguistic tree of 20 languages). Furthermore, it is worth noticing that the both the PCA and the F_ST_ results suggest that a few geographically neighbouring linguistic groups from different sub-regions are genetically quite similar, e.g. Benabena in the North region but which also has low F_ST_ values to groups in the North-West region and Kamano in the East region which has low F_ST_ values also to the Mid region.

Next, to look further into the population structure in the region, we phased the data using SHAPEIT and then used ChromoPainter (CP) to infer the genome-wide proportion of DNA for which each of 320 EPHNG individuals shares a most recent ancestor with each of the other 319 individuals. Visual inspection of these proportions (Supplementary Figure 3) confirms the structure inferred by PCA and Fst, e.g. with increasing heterogeneity between linguistic groups along the North-West to South-East cline.

**Figure 3.**
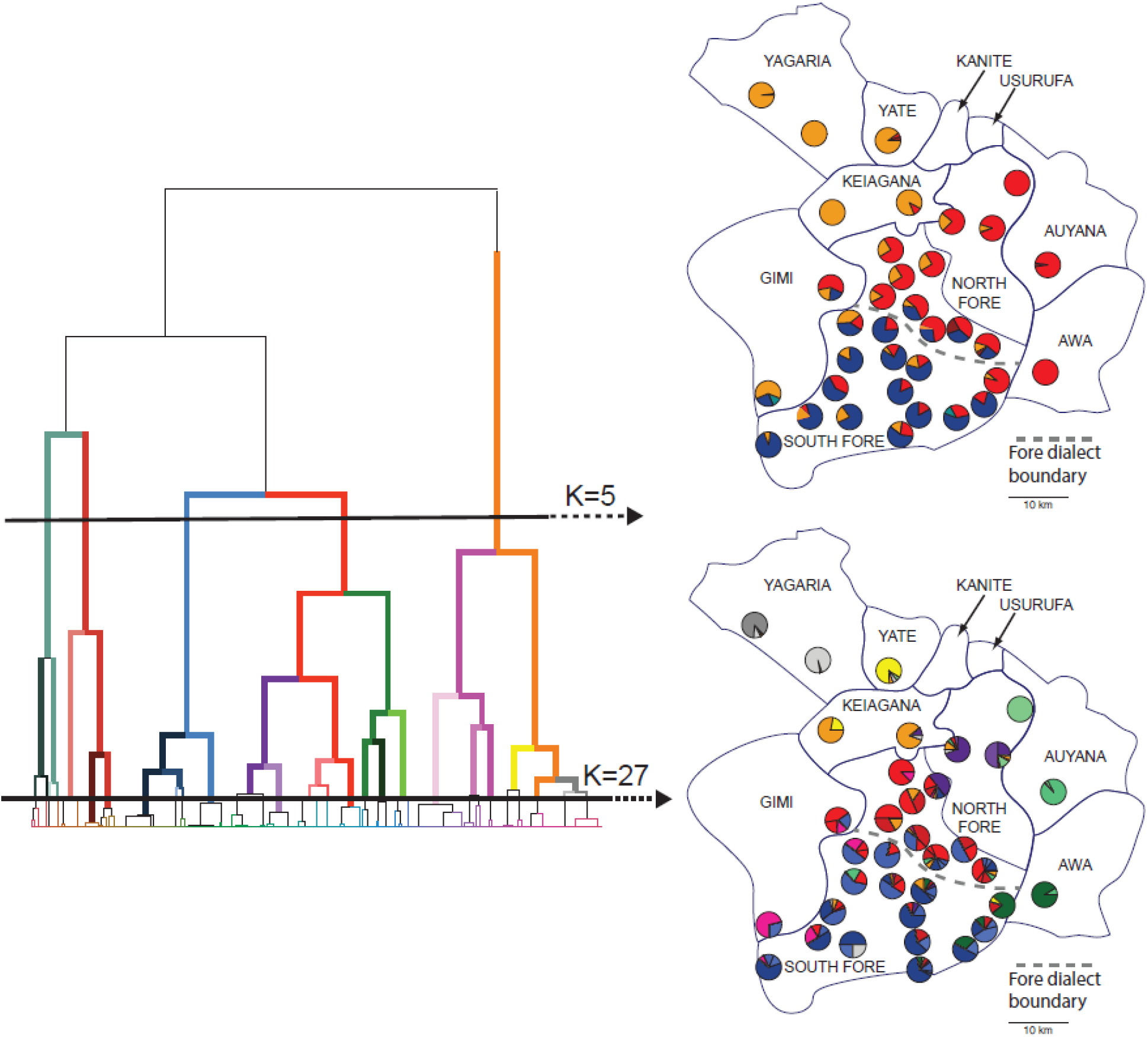
FS clustering output taken from analysis of 943 EHPNG individuals from 21 linguistic groups. a) FS dendogram with branches highlighted to match b) clustering of individuals for village sampling locations when K=5 and c) K=27

We next used FineStructure (FS) to assign individuals into discrete clusters based on the patterns of recent ancestry sharing inferred by CP. While the maximum posterior sample from FS assigned the 320 individuals into 38 clusters, FS then merges these clusters sequentially to generate a bifurcating tree (Figure 1a). At the level of the FS tree with 13 groups, individuals fall into groups broadly consistent with the patterns observed in F_ST_ and PCA. Moreover, several individuals cluster outside of their own linguistic group at this level of the tree, in a manner consistent with the outliers observed in PCA and supporting evidence of recent migration within the region.

To further investigate genetic differences between linguistic groups, we measured heterozygosity, runs of homozygosity (ROH) and linkage disequilibrium (LD) within each of the 20 EHPNG linguistic groups in the *Linguistic Group dataset* (Supplementary Table 2). All groups were downsampled to 13 individuals to obtain the same number of samples from each linguistic group after removal of the identified outliers/migrants. Concurring with the increasing genetic heterogeneity observed in the F_ST_ analysis, the trends in both average heterozygosity and average number of ROHs per linguistic group show a reduction in genetic diversity when moving through the region from North-West to South East. The Anga, Pawaian and Tairora have markedly higher average number of ROHs than the other groups which did not seem to be driven by differences in relatedness. Consistent with this, groups at the southern fringes, east and south east exhibit higher LD (Figure 2). Notably, LD seems to decay over physical distance much slower in the EHPNG groups than in CEU or YRI samples, likely reflecting the region’s history of isolation and small population sizes.

### Analysis of population structure at ‘FineScale’

Previous analyses using the *Linguistic Group dataset* permitted investigation of the relevance of the linguistic group classification in defining population structure without concerns of sample size biases impacting inferences. However, the data collection has an additional 623 samples largely from multiple villages within the North and South Fore groups allowing us to explore population structure at a finer-scale. This *Village Analysis dataset* contained 940 individuals from same 20 EHPNG linguistic groups plus 3 individuals from a 21^st^ linguistic group; the Kanite.

When we analysed the CP output for the subset of 943 EHPNG individuals using FS and, like earlier, clustered into 13 groups based on ascending the FS tree, the resulting groupings again largely mirror the linguistic group analysis (Supplementary Figure 4). However, notably there appear to be some clear fragmentation within several of the groups, especially the heavily sampled South Fore linguistic group (See Supplementary Table 3). Furthermore, when we stratified the FS analysis of the *Village Analysis dataset* by village instead of linguistic groups, the results showed clear evidence of population structure at village level. This is best encapsulated at 2 junctures in the FS tree (Figure 3). At FS K=5 (when the FS tree is ascended so that individuals are placed in 5 discrete clusters) there is the appearance of a split that largely fragments individuals within the South Fore linguistic group. At a village level this divergence clearly correlates with village of residence in terms of the Fore dialect being spoken. Individuals in villages speaking the Pamasugina dialect predominantly belong to cluster 3 and those speaking the Atigina and Ibusa dialects to cluster 4. Furthermore, at K=27 there is an observable pattern of fragmentation that corresponds to village of residence. This pattern largely reflects linguistic group membership with some aberrations. Individuals from the South Fore village of Ilesa, which is across the linguistic frontier from the Awa linguistic group show a clustering pattern more akin to the Awa sampled village than to other South Fore villages. Similarly, individuals in the village Kalu in the North Fore appear more genetically related to the Auyana than to the other two North Fore villages. There is a clear clustering pattern in the two sampled Yagaria villages at this level and further afield the two Anga sampled villages have completely distinct clustering patterns.

**Figure 4.**
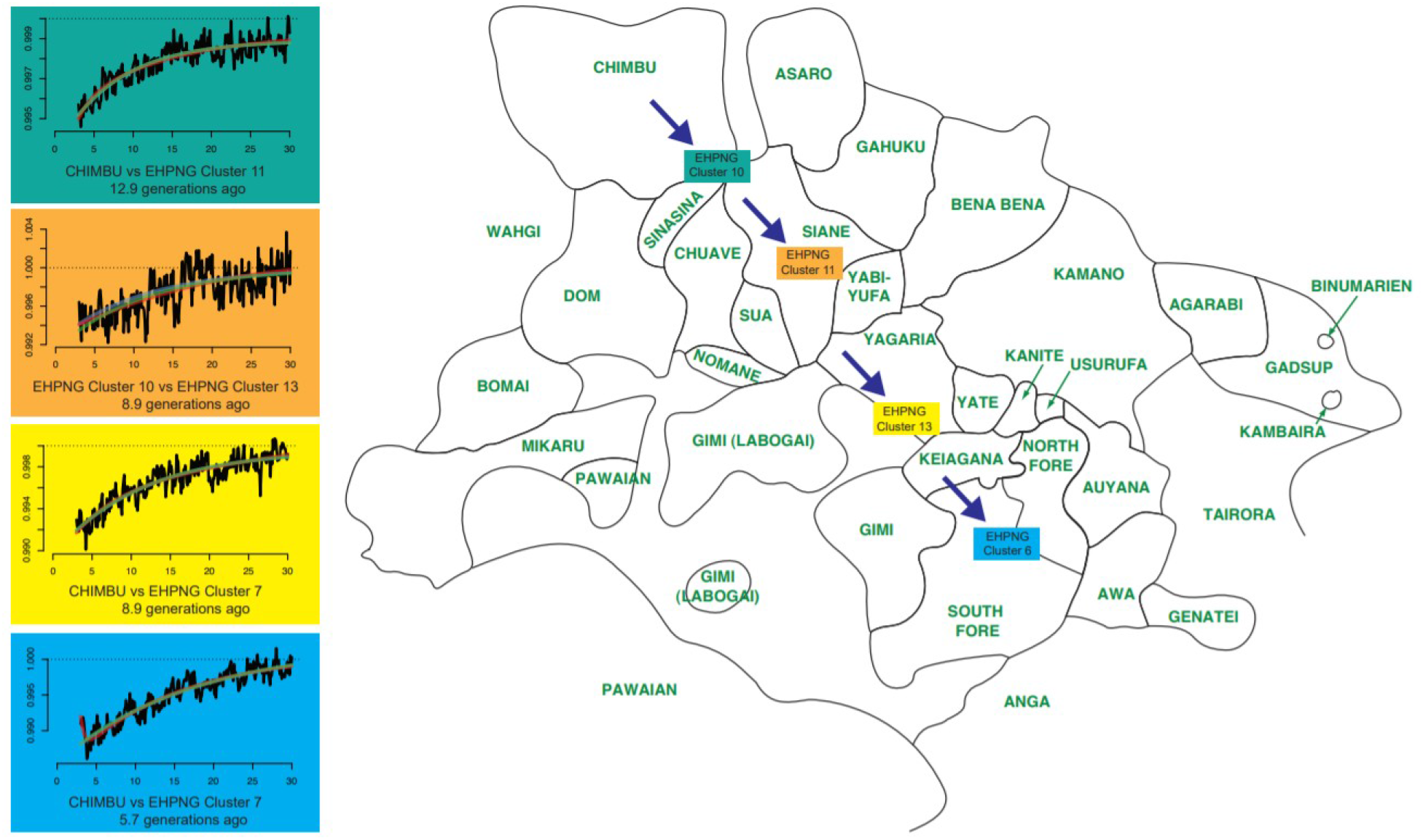
GLOBETROTTER results showing successive admixture events. Admixture pulses detected in a) *EHPNG Cluster 10* (Northwest) b) *EHPNG Cluster 11* (BenaBena/Labogai) c) *EHPNG Cluster 13* (Yagaria/Yate/Keiagana). Each admixture episode highlighted shows estimated date of admixture and accompanying admixture curves.

### Relationship of EHPNG populations to other groups

As EHPNG groups came into contact with Europeans in the 20^th^ century for the first time and contact with Europeans has led to admixture many other places in the world, we investigated if there are signs of admixture between EHPNG and Europeans using ADMIXTURE. Specifically, we performed an ADMIXTURE analysis for each linguistic group by extracting the 16 not closely related individuals from the group also present in the *Linguistic Group Dataset* as well as all the 1000 genomes populations CEU (North Americans with European ancestry) and CHB (Chinese individuals from Beijing) individuals and applied ADMIXTURE assuming the presence of 3 ancestral populations (K=3). These analyses revealed no individuals with significant CEU admixture proportions amongst the EHPNG, but four individuals displayed minor CHB admixture proportions (Supplementary Figure 5).

We also explored the relationship between EHPNG to other PNG populations by performing a CP analysis using the 943 EHPNG individuals combined with publicly available data from 353 individuals from other PNG regions, the *External Analysis* dataset (Supplementary Table 1 for sample description). Individuals were grouped into the 13 FS derived population clusters from the *Village Analysis dataset* (Supplementary table 3) which we believed best describes the overall population structure of the region. Visual inspection of the CP heatmap (Supplementary Figure 6) shows differential patterns of copying between the different *EHPNG* clusters in relation to non EHPNG populations. There appears to be marked affinity between *EHPNG Cluster 10 individuals* (mainly comprised of individuals from the Northwest of EHPNG) and a cluster formed of individuals from the neighbouring Chimbu population, which is administratively in a region neighbouring EHPNG to the west. Also, *EHPNG Cluster 1* (mainly comprised of Anga individuals) and *EHPNG Cluster 2* (mainly comprised of Pawaian individuals), have markedly different patterns of copying in this analysis compared to other *EHPNG Clusters* and to each other. When FS clustering was performed, individuals from *EHPNG Cluster 10* (Northwest) were placed closer in the tree alongside Chimbu individuals than to other EHPNG individuals. Individuals from *EHPNG Cluster 1* (Anga) and 2 (Pawaian) are placed at the extreme end of this tree (Supplementary Figure 6). 11 Individuals in the ‘Fine scale’ CP and FS analysis who appeared as recent long-distance migrants from other EHPNG groups clustered with other Highland populations entirely, possibly reflecting inter-region migration. All of these individuals were born in the 1960’s or later (after European contact) and included 2 individual outliers identified in PCA (Figure 1b).

To control for factors such as unequal sample size and incomplete lineage sorting that CHROMOPAINTER does not directly address, we used SOURCEFIND to infer the proportion of DNA for which each of the 13 EHPNG clusters shares most recent ancestry with a set of other “surrogate” groups mean to represent ancestral sources of the EHPNG,. For these surrogate groups, we used FS to cluster individuals from non-EHPNG populations into 13 genetically homogeneous groups (see Materials and Methods for details). We then applied SOURCEFIND to each of the 13 *EHPNG Clusters* using these 13 non-EHPNG clusters and the other 12 *EHPNG Clusters* as surrogates. Populations that did not contribute at least 5% to the ancestry profile of a single *EHPNG cluster* were excluded. Three of the 13 *EHPNG Clusters*, clusters 1 (Anga), 2 (Pawaian) and 10 (Northwest) have distinct distributions of inferred ancestry proportions (Supplementary Table 4) from the remaining ten clusters, which have profiles comprised of clusters that are nearest neighbours geographically within EHPNG. *EHPNG cluster 1* (Anga), has a distinct band of inferred ancestry from the Gulf Akoye, which is an Angan speaking group outside EHPNG that live close to the southern coast. *EHPNG Cluster 2* (Pawaian) finds the Southern Kiwai population as its majority contributory source. This population is located over 300km away outside EHPNG on the southern coast. *EHPNG cluster 10* (Northwest) has a strong contribution from the neighbouring Chimbu.

A second analysis which permitted only external non-EHPNG populations to act as sources showed that 8 of the 13 *EHPNG Cluster* populations has identical external copying (to the Southern Kiwai population), but *EHPNG Cluster 11* (BenaBena and Labogai) is a near equal mixture of this Southern Kiwai component and the Chimbu population neighbouring the region to the North-west. Moreover, this Chimbu component is close to 100% in *EHPNG Cluster 10* (Northwest) in this analysis. *EHPNG Cluster 9* (Awa) has a strong Gulf Akoye component in addition to the Suthern Kiwai component, this may represent some historic gene flow with the neighbouring Anga population to the south which was not visible when other populations were available as sources in the first analysis. The Anga and Pawaian maintain the same ancestry source composition as the first analysis.

To clarify whether some of these signals are due to recent admixture, We next applied GLOBETROTTER (GT) to infer admixture events in each of the 13 EHPNG clusters, using all other EHPNG and non-EHPNG clusters (as defined in the SF analysis above) as potential surrogates to the (unknown) admixing source groups (Supplementary Table 5 for summary of GT results). Multiple EHPNG clusters showed evidence of intermixing with their nearest neighbours, with inferred dates increasingly more recent going from the Northwest to the Southwest of EHPNG (Figure 4). In particular the earliest inferred date of admixture was in *EHPNG Cluster 10* (Northwest) at 12.8 generations ago, and the most recent date in *EHPNG Cluster 6* (North Fore) at 5.7 generations. This trend in dates appears to match the timescale and order of uptake of the sweet potato in the region(32). GT did not find any signals for *EHPNG Clusters 1* (Anga) and *2* (Pawaian) that would explain the differential SF ancestry profiles, suggesting that the distinct ancestry profiles of these groups compared to other EHPNG groups are more ancient in origin and not a result of recent admixture.

### Investigation of kuru-influenced sex biased migration informed by population structure understanding

As noted earlier, the presence of long distance migration was detectable due to aberrant clustering of individuals in FS and placement in PCA. Historically, kuru primarily affected only a few linguistic groups, including the South Fore, leaving some villages almost depleted of adult women. Therefore, we investigated if the kuru epidemic affected the sex ratio of migration into the South Fore group toward more females. To do so we identified migrants among the 331 individuals in the most affected South Fore linguistic group. Migrants into the group were assigned based the previously presented FS analyses of the *Village Analysis dataset* while taking into account our fine-scale structure results (for details see Methods). This approach resulted in 52 individuals in the South Fore group being classified as migrants out of 331 in total, of which 44 were female and 8 male, whereas the number of females and male among the non-migrants were 172 and 105, indicating an excess of females among the migrants (Chi squared test, p = 0.0017). For comparison, we performed the same analysis of the 376 individuals from 15 non-kuru affected linguistic groups. Here 44 individuals were classified as migrants of which 22 are females and 22 are males, whereas the number of females and males among the non-migrants are 167 and 165, suggesting no similar bias in these groups (Chi squared test, p = 0.97).

## Discussion

We have presented an in-depth study of the population structure of a remote highland region of PNG, based on an entirely new dataset that includes most of the relevant linguistic groups and extensive sampling by village, affording a level of resolution that has not previously been possible. We show that despite only covering an area of 11,157 km^2^, roughly the size of the island of Jamaica, the genetic differentiation between linguistic groups in the EHPNG region is strikingly high (maximum F_ST_ = 0.064, average F_ST_ = 0.024, average distance 45km). This differentiation is comparable to that previously reported for the entire PNG Highland region(17), of which EHPNG is only a small sub-region. Furthermore, our analyses have revealed the presence of complex population structure even at dialect and village level. Whilst of interest for understanding the origins of modern populations, these findings also provide the background for study of the genetic impact of the large-scale prion disease epidemic, kuru.

A key question is what factors have contributed to such strong population structure in such a small region. One possible factor is that while the Western Highlands had optimal conditions for Taro cultivation, which originated there, the conditions were suboptimal in EHPNG(33). Indeed, some groups in EHPNG have been described as having been ‘proto-agriculturalists’ retaining elements of hunter-gatherer subsistence in their lifestyles(33). Analogous to other H-G groups, most EHPNG linguistic groups have greatly reduced population densities compared to other highland regions(1) and lower historical effective population sizes(17). These reduced population densities would likely have led to increased effects of genetic drift between groups.

In addition to this, discrete historical events, such as the arrival of goods like dogs(34), pigs(35) and chickens to the region, climatic changes and epidemics are also likely to have played a role in shaping population structure in the region. This is supported by the detection of admixture dates that correlate strongly with the arrival and estimated uptake of the sweet potato by different groups in the region.

In addition to geography, overall broad-scale population structure correlates with linguistics. For example, the best estimated linguistic tree for these groups analysed shows the Pawaian and Anga as outgroups (Figure 1c), echoing the genetic analysis. We even see examples of fine scale parallels between linguistic and genetic differences in the Fore dialect groups. However, the correlation between linguistic groupings and geographical regions it is difficult to disentangle the relative role of these two factors.

The simple colonial era definition of linguistic groups, which has been used in previous genetic studies, does not always satisfactorily describe population structure in the region. For example, linguistic groups are barely distinguishable from one another genetically in the north-west of EHPNG. This relative homogeneity may be due to more intensely practiced agriculture and higher population densities in this sub-region(10). It may also be attributable to shared recent admixture events we detect that may be related to genetic inflow from the neighbouring Chimbu region to the west of EHPNG. Furthermore, in the village level analyses of the Fore, we found genetic affinity of some villages to be closer to non-Fore groups. This observation is not completely surprising, as it is well understood that the clan structure of political, economic and social unions that comprised the precolonial landscape in EHPNG often spanned linguistic group boundaries. This observed signature (for example as observed with Ilesa in the South Fore, Figure 3) could also be the result of a village founding event when a whole village is uprooted (e.g. due to conflict) and moves considerable distance to new territory, with members acquiring the language of their new residence. Such founding events have been observed in the anthropological record before within EHPNG(11).

Several analyses revealed the presence of recent long-distance migration in the region. For instance, we found three potential migrants from the Fore linguistic group into the Pawaian (whose sample records showed their presence consequent upon a woman marrying into the Pawaian and taking two family members with her). However, marriages across the Fore/Pawaian divide are believed to be very rare if they ever occurred in pre European contact times, due to the considerable barriers of endemic warfare and extreme terrain. Consistent with this we found that all three observed migrants were born after European contact, which resulted in a cessation of warfare and the development of transport infrastructure which may have facilitated these movements. Hence it is possible that such long distance migration is a recent phenomenon, consistent with the two groups being so genetically distant.

A final example demonstrating that analysis of population structure using linguistic group labels is not fully satisfactory was where we observed a clear resolution of village differences based on South Fore dialect spoken within the linguistic group. This suggests that pooling all South Fore into a single population may not adequately capture population structure. Furthermore, it reveals the dynamic and ongoing processes of cultural and demographic change that has been unfolding in the region. Given the small genetic differences between the FS clusters that represent the two dialect groups, the small geographic differences between different dialect villages and the fact that marital exchange and migration was known to occur between villages across the dialect divide it is likely such a split was recent in origin. This echoes oral origin histories held amongst the Fore which details the expansion and fragmentation of the Fore people into the three distinct dialect groups(6).

One of the population structure patterns that very clearly follows the linguistic groupings is the fact that the Anga and Pawaian linguistic groups appear not only highly distinct from all other groups, but also from each other. Clusters comprising these groups had highest genetic similarity with groups outside of EHPNG, rather than EHPNG neighbours as was the case in all other clusters (except *EHPNG Cluster* 10 who had a strong contribution with the neighbouring Chimbu). In the case of the Pawaian, the inferred closest ancestry source was the Southern Kiwai, a coastal population over 300km from EHPNG. Interestingly, the Pawaian linguistic group is known to live semi-nomadically in forests at lower elevations and lower population densities than the rest of the region(1), which may in itself explain why they have ended up somewhat genetically distinct from the rest of the EHPNG groups. The oral histories held by the Pawaian speak of long migrations through uninhabited forest regions originating from coastal regions(23).

Our results do not support significant genetic influence from the outside of the PNG highlands. In particular, unlike in many other regions of the world that have been colonized by countries with people of European descent(30), we found no signs of European ancestry in EHPNG individuals (Supplementary Figure 5). And while we did in our ADMXITURE analyses observe some signatures that are consistent with some admixture with people of East Asian ancestry, this could just as well be caused by other PNG populations not being represented in those analyses (17). When compared to other PNG populations external to EHPNG, 8 of the 13 population clusters have near identical external ancestry profiles despite being genetically distinct from each other in other analyses. No recent admixture was detected with external groups for these 8 population clusters, suggesting that genetic differences between them have arisen since any shared external relationship and driven by group separation and genetic drift. This supports previously suggested population histories(1, 17) with an expansion of groups (‘neolithic expansion’) emanating from the Western Highlands as a result of the development of Taro agriculture and displacement of previous groups that lived there, possibly ancestral to the Anga who have greatly distinct ancestry profiles in our analyses and that now live in the southern fringes of the region. In fact ‘Anga-like’ artefacts, believed to be ancient have been found as far north as in the Kamano linguistic group, reflecting a more widely dispersed settlement in the region(1).We additionally find evidence of more recent dispersals of peoples emanating from the west possibly due to the arrival and impact of the sweet potato.

The observation of recent migrants throughout analysis allowed the discovery of significant sex-biased migration into the South Fore linguistic group. This observation is consistent with accounts from the region that notes that during the epidemic the near absence of adult women in villages with high kuru incidence(21). Men would frequently marry multiple times as a result of their previous wives dying form kuru, and strains were also placed on communities as a result of increased child care burden(9). Hence the observed excessive inflow of female recent migrants into South Fore villages may have been the result of the need to replace lost adult women and mitigate the impact of kuru on Fore society. The smaller amount of male migration could also have been due to fears of sorcery for which the Fore had a fearsome reputation(36). Kuru was linked to sorcery amongst communities in EHPNG. With deaths from kuru rising to be the leading cause of death in the mid-20^th^ century within the South Fore, associated fears of sorcery are likely to have risen as well.

In summary, our results suggest that the observed population structure is not driven by admixture from outside the highland PNG region, which is consistent with the historical record of EHPNG region being isolated until recently. Also, while the population structure does to some extent mimic the linguistic groupings in the area, we observe several patterns of population structure that suggest that the different linguistic groups are not entirely genetically distinct and isolated from each other. This is consistent with previous knowledge of clans playing a key role in the cultural grouping in the area and of the presence of cultural features that aided possible migration between neighbouring linguistic groups in the region. Finally, we observed signs that long distance migration has at least in more recent times taken place. Importantly this, in combination with the understanding of population structure, has permitted a novel analysis of sex-biased flows of migration that are likely to have been impacted by kuru. This highlights the essential nature of understanding population structure of a region prior to attempting to investigate hypotheses regarding the impact of epidemics on affected populations.

The population structure of EHPNG reveals a complex multi-layered set of factors that have caused high population differentiation, likely including both geographic and cultural factors. Furthermore, it suggests that the current population structure may still be evolving, at least if the migration between distant parts of the region is indeed a new phenomenon as our results suggest. Overall our results demonstrate that simplistic descriptions of the population structure in regions like EHPNG based on linguistic groupings presumed to be static, are likely to neglect the far richer texture of dynamic forces and history that has shaped communities.

## Acknowledgments

We acknowledge the help of the Eastern Highlands Province communities and the many individuals who have made this study possible. We thank the Director of the Papua New Guinea Institute of Medical Research, Professor William Pomat, and the former Directors, Professor John Reeder and Professor Peter Siba, for their support. We also thank the kuru project field staff whose hard work made this study possible.

