## SUPPLEMENTARY MATERIAL for "Population Structure and Migration in the Eastern Highlands of Papua New Guinea; a Region Impacted by the kuru Epidemic"

**Supplementary Figure 1**

**Principal Components 1-10 of 320 individuals from 20 EHPNG linguistic groups**

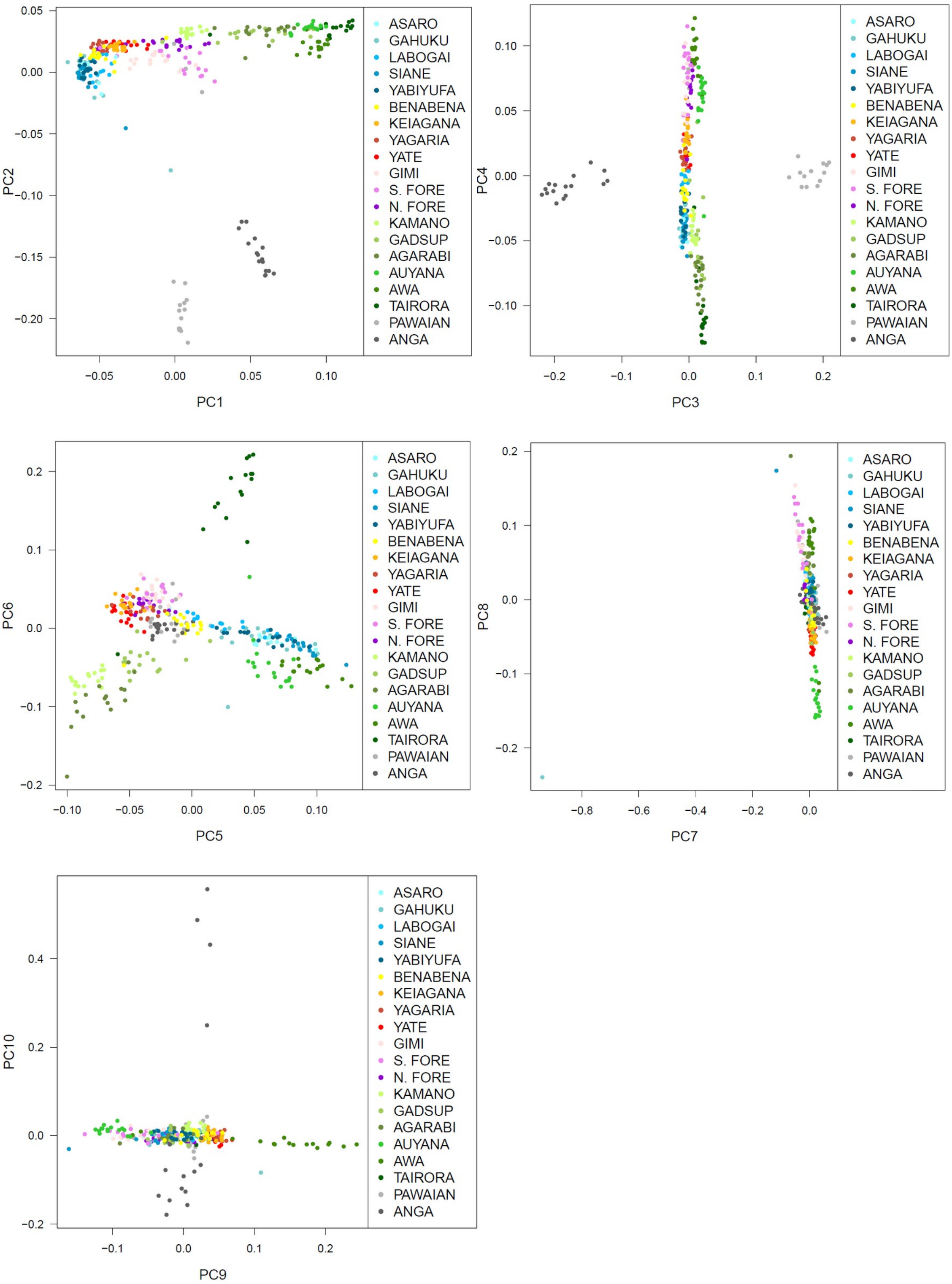

**Supplementary Figure 2 – FST analysis of 20 EHPNG linguistic groups after 7 outlier individuals identified in PCA were removed**

| Sub-Region | North-West | | | | | North | | | | Mid | | | East | | | South-East | | Boundary Groups | | |
|---|---|---|---|---|---|---|---|---|---|---|---|---|---|---|---|---|---|---|---|---|
| Group | ASARO | GAHUKU | SIANE | YABIYUFA | LABOGAI | BENABENA | YAGARIA | YATE | KEIAGANA | NF | GIMI | SF | KAMANO | AGARABI | GADSUP | AWA | AUYANA | TAIRORA | PAWAIAN | ANGA |
| ASARO | N/A | 0.00208813 | 0.00323946 | 0.00304656 | 0.00482923 | 0.00450205 | 0.0100217 | 0.0112801 | 0.0102595 | 0.0139273 | 0.0130712 | 0.0136509 | 0.0177209 | 0.0248563 | 0.0213094 | 0.033871 | 0.0321877 | 0.0364874 | 0.0447713 | 0.0427318 |
| GAHUKU | 0.00208813 | N/A | 0.00403269 | 0.00307137 | 0.00442292 | 0.00402141 | 0.00878225 | 0.0108087 | 0.00960807 | 0.014016 | 0.0126661 | 0.0131946 | 0.0177256 | 0.0248036 | 0.0211576 | 0.033845 | 0.0325416 | 0.0365621 | 0.04443 | 0.0426309 |
| SIANE | 0.00323946 | 0.00403269 | N/A | 0.00295873 | 0.00565179 | 0.00794308 | 0.0130397 | 0.0144156 | 0.0133232 | 0.0169674 | 0.015522 | 0.0163432 | 0.0213885 | 0.0279519 | 0.023913 | 0.0365545 | 0.0347022 | 0.039508 | 0.0459839 | 0.0447046 |
| YABIYUFA | 0.00304656 | 0.00307137 | 0.00295873 | N/A | 0.00317057 | 0.00422825 | 0.00876864 | 0.0107637 | 0.00944112 | 0.013341 | 0.0120313 | 0.0129408 | 0.0175335 | 0.0250269 | 0.0211325 | 0.0338617 | 0.0321979 | 0.0367466 | 0.0445344 | 0.0423319 |
| LABOGAI | 0.00482923 | 0.00442292 | 0.00565179 | 0.00317057 | N/A | 0.00352929 | 0.00765867 | 0.00940833 | 0.00825359 | 0.0115451 | 0.00958202 | 0.0110181 | 0.0166088 | 0.0238669 | 0.0196181 | 0.0322544 | 0.0304812 | 0.0349638 | 0.0434166 | 0.0412599 |
| BENABENA | 0.00450205 | 0.00402141 | 0.00794308 | 0.00422825 | 0.00352929 | N/A | 0.0049936 | 0.00664453 | 0.00589182 | 0.0100517 | 0.00927711 | 0.0103766 | 0.0135266 | 0.0222891 | 0.0185133 | 0.0323639 | 0.0305942 | 0.0346303 | 0.0448317 | 0.0425984 |
| YAGARIA | 0.0100217 | 0.00878225 | 0.0130397 | 0.00876864 | 0.00765867 | 0.0049936 | N/A | 0.00693138 | 0.00422451 | 0.0108417 | 0.011201 | 0.0116966 | 0.0164067 | 0.0247372 | 0.0212483 | 0.0346149 | 0.0328665 | 0.0372314 | 0.0488231 | 0.0459137 |
| YATE | 0.0112801 | 0.0108087 | 0.0144156 | 0.0107637 | 0.00940833 | 0.00664453 | 0.00693138 | N/A | 0.0032287 | 0.00923029 | 0.0116523 | 0.0114735 | 0.0143852 | 0.02249 | 0.0190379 | 0.0326633 | 0.029878 | 0.0347369 | 0.0482078 | 0.0449578 |
| KEIAGANA | 0.0102595 | 0.00960807 | 0.0133232 | 0.00944112 | 0.00825359 | 0.00589182 | 0.00422451 | 0.0032287 | N/A | 0.00748904 | 0.00884676 | 0.00904965 | 0.0149186 | 0.0223274 | 0.0189801 | 0.031453 | 0.0293169 | 0.0342976 | 0.0466464 | 0.0439632 |
| NORTH FORE | 0.0139273 | 0.014016 | 0.0169674 | 0.013341 | 0.0115451 | 0.0100517 | 0.0108417 | 0.00923029 | 0.00748904 | N/A | 0.00834718 | 0.00578027 | 0.0157325 | 0.0215307 | 0.0169038 | 0.0256082 | 0.0233218 | 0.031258 | 0.0474791 | 0.0436238 |
| GIMI | 0.0130712 | 0.0126661 | 0.015522 | 0.0120313 | 0.00958202 | 0.00927711 | 0.011201 | 0.0116523 | 0.00884676 | 0.00834718 | N/A | 0.00244127 | 0.0169832 | 0.0224852 | 0.0182146 | 0.027171 | 0.0265544 | 0.0323622 | 0.0446674 | 0.0413095 |
| SOUTH FORE | 0.0136509 | 0.0131946 | 0.0163432 | 0.0129408 | 0.0110181 | 0.0103766 | 0.0116966 | 0.0114735 | 0.00904965 | 0.00578027 | 0.00244127 | N/A | 0.0166215 | 0.0206169 | 0.0160705 | 0.0235983 | 0.0234752 | 0.029906 | 0.0436877 | 0.0389558 |
| KAMANO | 0.0177209 | 0.0177256 | 0.0213885 | 0.0175335 | 0.0166088 | 0.0135266 | 0.0164067 | 0.0143852 | 0.0149186 | 0.0157325 | 0.0169832 | 0.0166215 | N/A | 0.0152002 | 0.0125893 | 0.0317508 | 0.0279548 | 0.0316636 | 0.0518332 | 0.0478602 |
| AGARABI | 0.0248563 | 0.0248036 | 0.0279519 | 0.0250269 | 0.0238669 | 0.0222891 | 0.0247372 | 0.02249 | 0.0223274 | 0.0215307 | 0.0224852 | 0.0206169 | 0.0152002 | N/A | 0.00872393 | 0.0281886 | 0.0255052 | 0.0262098 | 0.0545001 | 0.0489604 |
| GADSUP | 0.0213094 | 0.0211576 | 0.023913 | 0.0211325 | 0.0196181 | 0.0185133 | 0.0212483 | 0.0190379 | 0.0189801 | 0.0169038 | 0.0182146 | 0.0160705 | 0.0125893 | 0.00872393 | N/A | 0.0228497 | 0.0192087 | 0.019972 | 0.0501189 | 0.0453343 |
| AWA | 0.033871 | 0.033845 | 0.0365545 | 0.0338617 | 0.0322544 | 0.0323639 | 0.0346149 | 0.0326633 | 0.031453 | 0.0256082 | 0.027171 | 0.0235983 | 0.0317508 | 0.0281886 | 0.0228497 | N/A | 0.0197304 | 0.0322197 | 0.0595577 | 0.051597 |
| AUYANA | 0.0321877 | 0.0325416 | 0.0347022 | 0.0321979 | 0.0304812 | 0.0305942 | 0.0328665 | 0.029878 | 0.0293169 | 0.0233218 | 0.0265544 | 0.0234752 | 0.0279548 | 0.0255052 | 0.0192087 | 0.0197304 | N/A | 0.0287577 | 0.0598465 | 0.0536466 |
| TAIRORA | 0.0364874 | 0.0365621 | 0.039508 | 0.0367466 | 0.0349638 | 0.0346303 | 0.0372314 | 0.0347369 | 0.0342976 | 0.031258 | 0.0323622 | 0.029906 | 0.0316636 | 0.0262098 | 0.019972 | 0.0322197 | 0.0287577 | N/A | 0.0638242 | 0.056528 |
| PAWAIAN | 0.0447713 | 0.04443 | 0.0459839 | 0.0445344 | 0.0434166 | 0.0448317 | 0.0488231 | 0.0482078 | 0.0466464 | 0.0474791 | 0.0446674 | 0.0436877 | 0.0518332 | 0.0545001 | 0.0501189 | 0.0595577 | 0.0598465 | 0.0638242 | N/A | 0.0599953 |
| ANGA | 0.0427318 | 0.0426309 | 0.0447046 | 0.0423319 | 0.0412599 | 0.0425984 | 0.0459137 | 0.0449578 | 0.0439632 | 0.0436238 | 0.0413095 | 0.0389558 | 0.0478602 | 0.0489604 | 0.0453343 | 0.051597 | 0.0536466 | 0.056528 | 0.0599953 | N/A |

**Supplementary Figure 3 - CP heatmap of 320 individuals from 20 EHPNG linguistic groups**

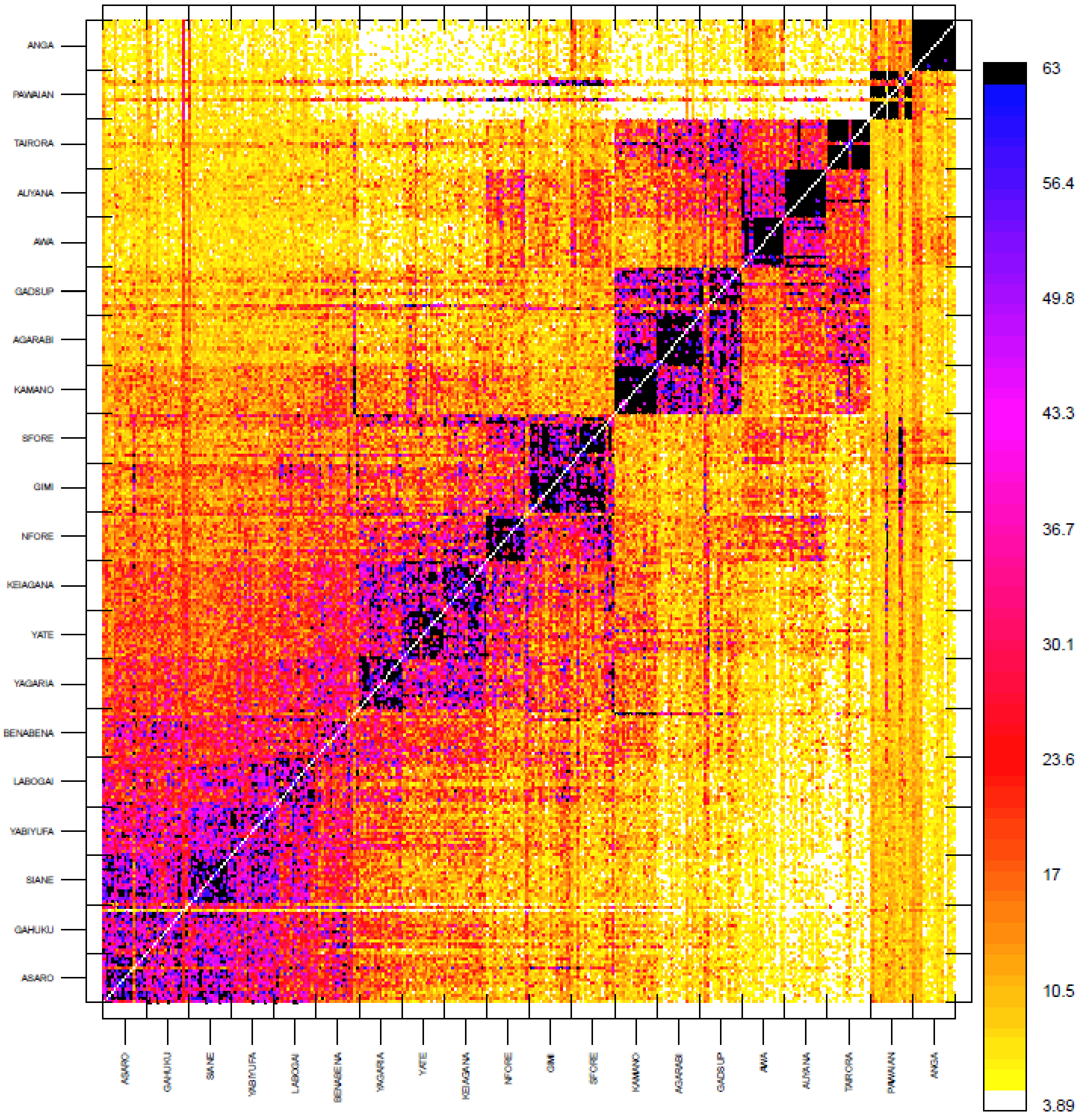

**Supplementary Figure 4 – a) CP heatmap of 943 individuals from 21 EHPNG Ethno-Linguistic Groups and plot of FS clustering for K=13grouped by 13 FS clusters. b) Same FS clustering but by linguistic group on EHPNG map**

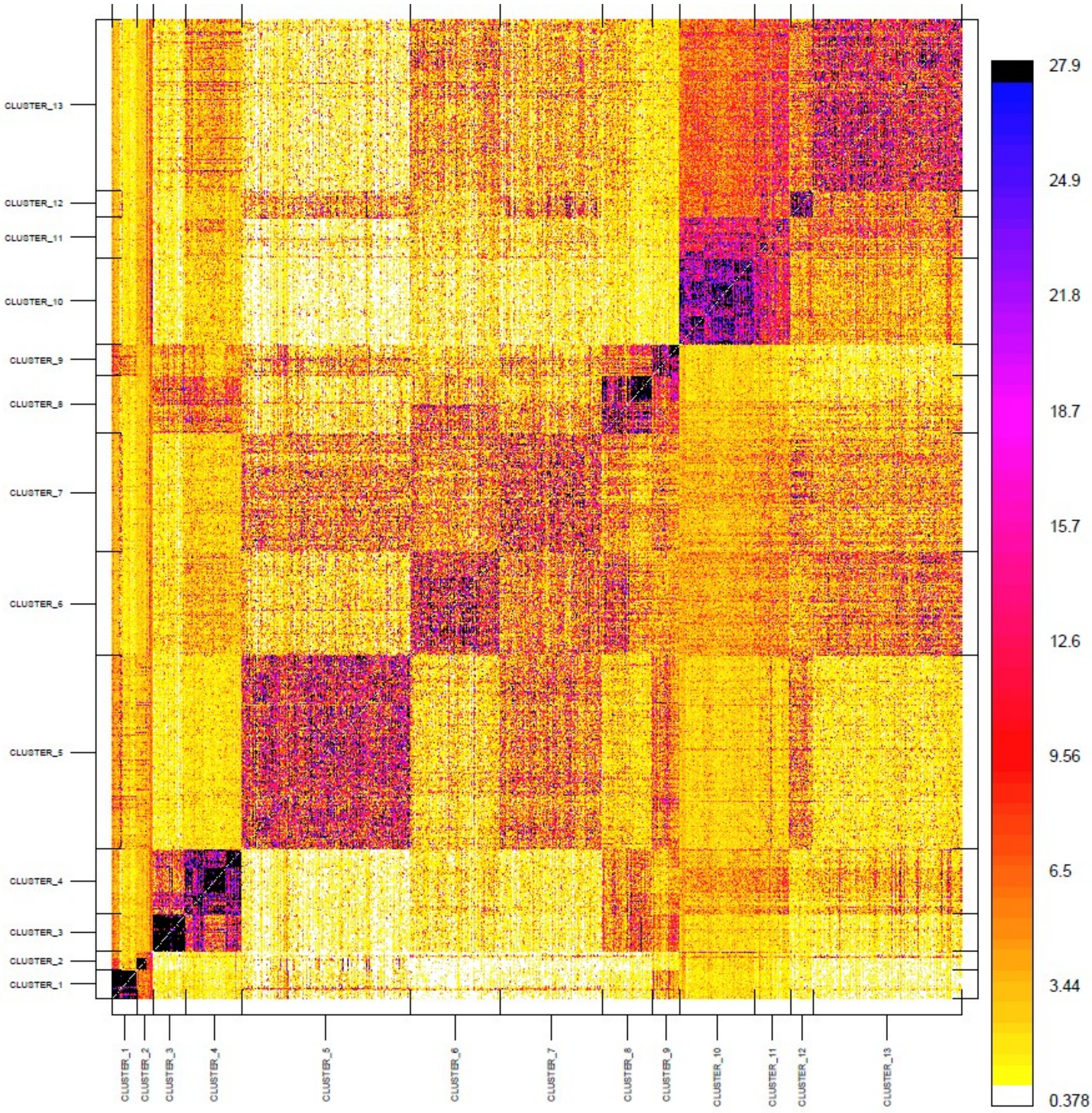
CLUSTER_1
CLUSTER_2
CLUSTER_3
CLUSTER_4
CLUSTER_5
CLUSTER_6
CLUSTER_7
CLUSTER_8
CLUSTER_9
CLUSTER_10
CLUSTER_11
CLUSTER_12
CLUSTER_13
27.9
24.9
21.8
18.7
15.7
12.6
9.56
6.5
3.44
0.378

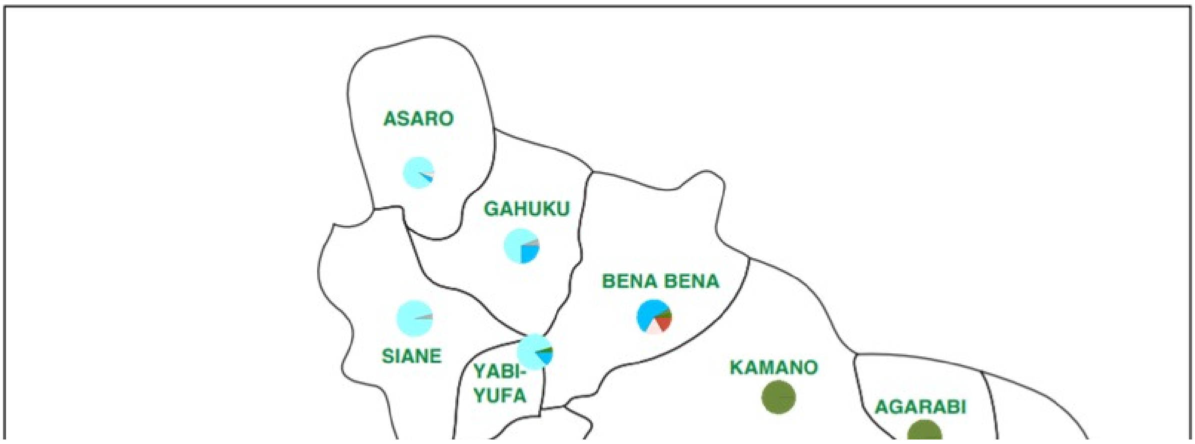
ASARO
GAHUKU
BENA BENA
SIANE
YABI-
YUFA
KAMANO
AGARABI

**Supplementary Figure 5 – ADMIXTURE analysis of EHPNG linguistic groups with 1000 genomes project populations CEU and YRI**

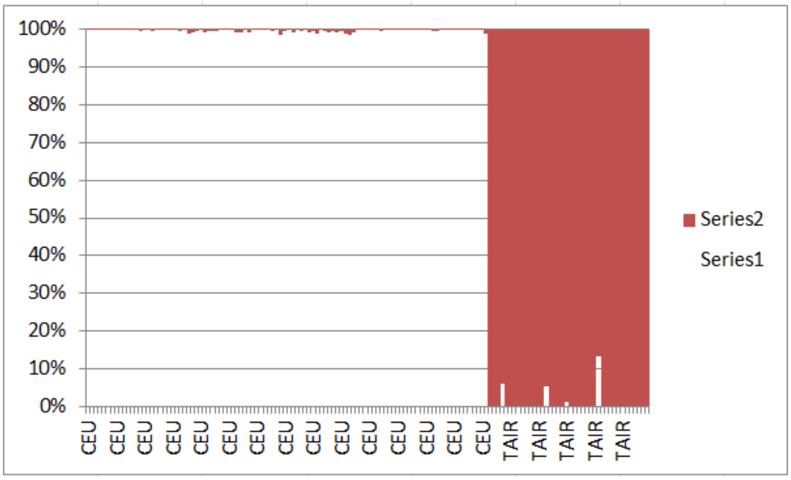


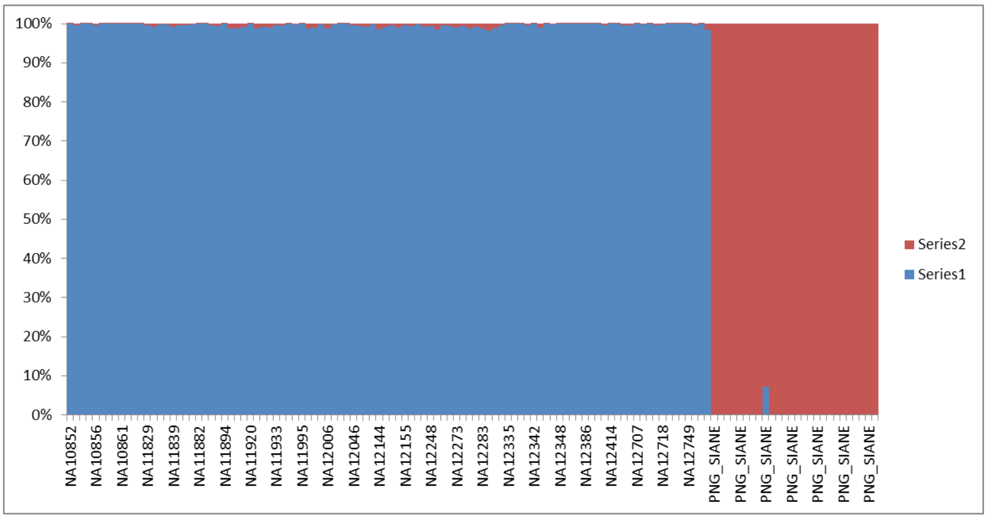


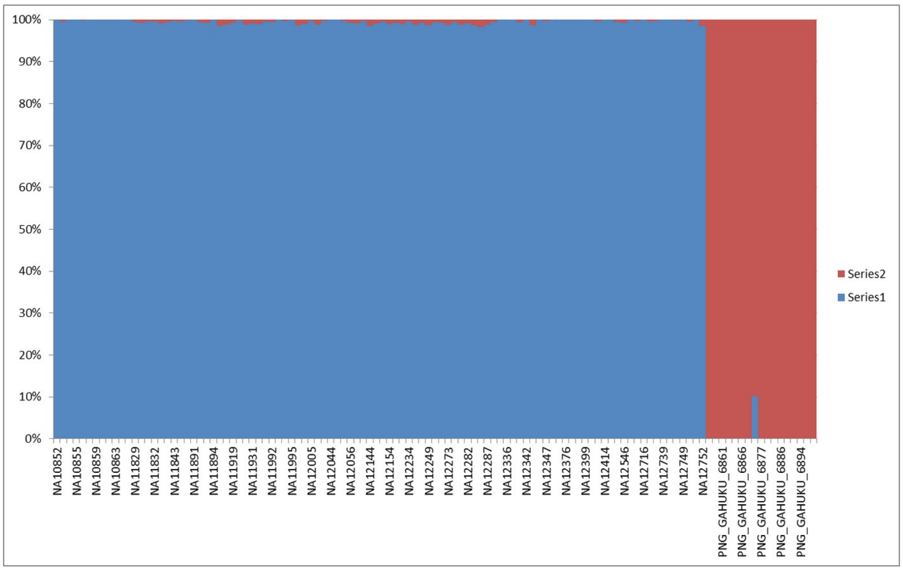

**Supplementary Figure 6 – a) CP heatmap of 1293 individuals, 943 from 21 EHPNG linguistic groups and 372 individuals from other PNG regions b) Accompanying FS tree**

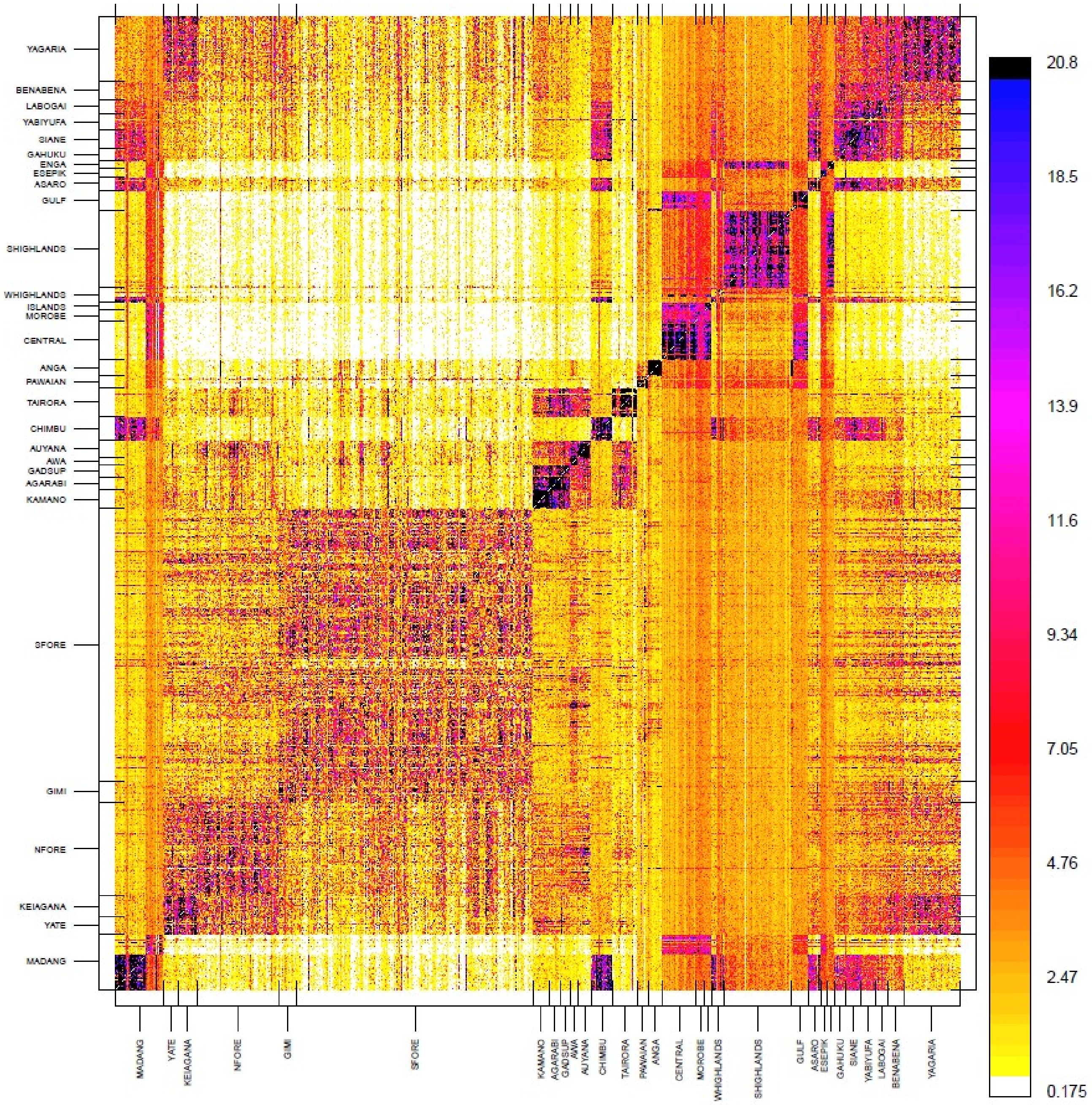

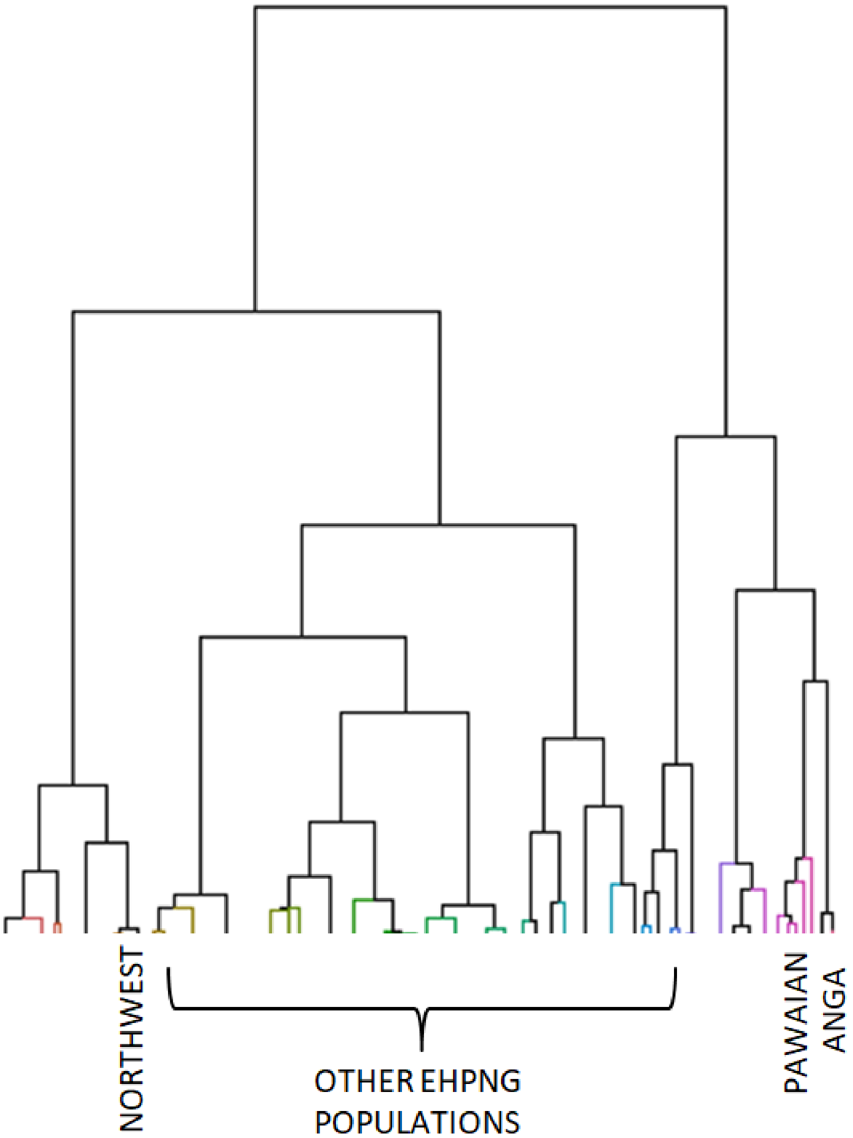
NORTHWEST
OTHER EHPNG
POPULATIONS
PAWAIAN
ANGA

**Supplementary Table 1 – Summary Data for individuals used in analyses**

**1a) Linguistic Group Breakdown of 'Linguistic Group' Dataset**

| **LINGUISTIC GROUP** | **COUNT OF INDIVIDUALS** |
|---|---|
| AGARABI | 16 |
| ANGA | 16 |
| ASARO | 16 |
| AUYANA | 16 |
| AWA | 16 |
| BENABENA | 16 |
| GADSUP | 16 |
| GAHUKU | 16 |
| GIMI | 16 |
| KAMANO | 16 |
| KEIAGANA | 16 |
| LABOGAI | 16 |
| NORTH FORE | 16 |
| PAWAIAN | 16 |
| SIANE | 16 |
| SOUTH FORE | 16 |
| TAIRORA | 16 |
| YABIYUFA | 16 |
| YAGARIA | 16 |
| YATE | 16 |
| **Grand Total** | **320** |

**Village breakdown of 'Village dataset'**

| **LINGUISTIC GROUP** / VILLAGE | COUNT OF INDIVIDUALS |
|---|---|
| **AGARABI** | |
| KAINOA | 17 |
| **ANGA** | |
| AMORA | 2 |
| BOIKO | 3 |
| DUNGWI | 15 |
| SIMBARI | 2 |
| **ASARO** | |
| GIMISEVI | 18 |
| **AUYANA** | |
| OIYANA | 22 |
| **AWA** | |
| TAUNA | 11 |
| **BENABENA** | |
| MATAUSA | 24 |
| **GADSUP** | |

| | |
|---|---|
| UKARUMPA | 16 |
| **GAHUKU** | |
| HORIPOKAVE | 16 |
| **GIMI** | |
| EMO | 15 |
| ETEVE | 8 |
| HAIYARU | 1 |
| UNKNOWN | 4 |
| **KAMANO** | |
| HOMORI | 24 |
| **KANITE** | |
| UNKNOWN | 3 |
| **KEIAGANA** | |
| HOGATERU | 18 |
| UWAMI | 9 |
| **LABOGAI** | |
| LOSAVE | 19 |
| **NORTHFORE** | |
| ANUMPA | 76 |
| AWANDE | 21 |
| KALU | 16 |
| OKAPA | 1 |
| UMASA | 1 |
| UNKNOWN | 11 |
| **PAWAIAN** | |
| UNKNOWN | 16 |
| **SIANE** | |
| WAIFO | 24 |
| **SOUTHFORE** | |
| KETABI | 1 |
| AGAKAMATASA | 12 |
| AI | 5 |
| AMORA | 1 |
| AWAROSA | 9 |
| HIGITARU | 1 |
| ILESA | 13 |
| INTAMATASA | 6 |
| IVAKI | 35 |
| KALU | 6 |
| KAMATA | 12 |
| KAMIRA | 16 |
| KANIGITASA | 23 |
| KASARAI | 3 |
| KEIAKASA | 2 |
| KETABI | 12 |
| KUME | 6 |
| MENTILASA | 9 |
| MIARASA | 9 |

| | |
|---|---|
| MUGAIAMUTI | 35 |
| OMA-KASORU | 3 |
| PAITI | 15 |
| PUROSA-TAKAI | 22 |
| TAKAI | 27 |
| TAKARI | 8 |
| TAMOGAVISA | 1 |
| UMASA | 5 |
| UNKNOWN | 23 |
| URAI | 4 |
| WAISA | 25 |
| WANIKANTO | 2 |
| WANITABI | 5 |
| YAGAREBA | 2 |
| YAGUSA | 1 |
| YASUBI | 4 |
| **TAIRORA** | |
| BONTA | 38 |
| **YABIYUFA** | |
| HOIHATOBE | 22 |
| **YAGARIA** | |
| KIWURUGA | 49 |
| NUSAGUNA | 38 |
| **YATE** | |
| KAUNA | 20 |
| Grand Total | 943 |

**1b) Summary of Additional Individuals Used in External Populations Analysis**

| PNG REGION | COUNT OF INDIVIDUALS |
|---|---|
| CENTRAL | 51 |
| CHIMBU | 32 |
| ENGA | 11 |
| ESEPIK | 11 |
| GULF | 27 |
| ISLANDS | 10 |
| MADANG | 74 |
| MOROBE | 14 |
| SHIGHLANDS | 102 |
| WHIGHLANDS | 20 |

**1c) Samples included in haplotype phasing panel**

| POPULAITON | NUMBER OF SAMPLES | ORIGIN |
|---|---|---|
| EHPNG | 1374 | PRIMARY DATA |
| PNG | 380 | ACCESS AGREEMENT |
| ACB | 95 | 1000 GENOMES |
| Altai | 1 | ANCIENT HOMININ |
| ASW | 45 | 1000 GENOMES |
| Batwa | 5 | AFRICA |
| BEB | 83 | 1000 GENOMES |
| CDX | 82 | 1000 GENOMES |
| CEU | 91 | 1000 GENOMES |
| CHB | 103 | 1000 GENOMES |
| CHS | 97 | 1000 GENOMES |
| CLM | 93 | 1000 GENOMES |
| Colla | 22 | SOUTH AMERICA |
| Denisova | 1 | ANCIENT HOMININ |
| ESN | 95 | 1000 GENOMES |
| FIN | 99 | 1000 GENOMES |
| GB20 | 1 | ANCIENT EUROPEAN |
| GBR | 85 | 1000 GENOMES |
| GIH | 96 | 1000 GENOMES |
| GWD | 112 | 1000 GENOMES |
| IBS | 107 | 1000 GENOMES |
| ITU | 98 | 1000 GENOMES |
| JPT | 104 | 1000 GENOMES |
| Khoesan_ColouredColesberg | 20 | AFRICA |
| Khoesan_ColouredWellingto | 20 | AFRICA |

| | | |
|---|---|---|
| n | | |
| Khoesan_GuiGhanaKgal | 15 | AFRICA |
| Khoesan_Juhoansi | 18 | AFRICA |
| Khoesan_Karretjie | 20 | AFRICA |
| Khoesan_Khomani | 39 | AFRICA |
| Khoesan_Khwe | 17 | AFRICA |
| Khoesan_Nama | 20 | AFRICA |
| Khoesan_SEBantu | 20 | AFRICA |
| Khoesan_SWBantu | 12 | AFRICA |
| Khoesan_Xun | 19 | AFRICA |
| KHV | 98 | 1000 GENOMES |
| Kuba_Bindi | 11 | AFRICA |
| Kuba_Dekese | 5 | AFRICA |
| Kuba_Dinga | 1 | AFRICA |
| Kuba_Kete | 12 | AFRICA |
| Kuba_Kuba | 24 | AFRICA |
| Kuba_Lele | 19 | AFRICA |
| Kuba_Luba | 7 | AFRICA |
| Kuba_Lubakat | 1 | AFRICA |
| Kuba_Luluwa | 47 | AFRICA |
| Kuba_Luntu | 11 | AFRICA |
| Kuba_Mbala | 1 | AFRICA |
| Kuba_Songe | 2 | AFRICA |
| Kuba_Tetela | 5 | AFRICA |
| Kuba_Tshokwe | 1 | AFRICA |
| LBK | 1 | ANCIENT EUROPEAN |
| Loschbour | 1 | ANCIENT EUROPEAN |
| LWK | 79 | 1000 GENOMES |
| MSL | 69 | 1000 GENOMES |
| MXL | 55 | 1000 GENOMES |
| PEL | 76 | 1000 GENOMES |
| PJL | 86 | 1000 GENOMES |
| PUR | 104 | 1000 GENOMES |
| Pygmy_Baka_Cam | 56 | AFRICA |
| Pygmy_Baka_Gab | 16 | AFRICA |
| Pygmy_Bakiga | 34 | AFRICA |
| Pygmy_Batwa | 27 | AFRICA |
| Pygmy_Bongo_GabE | 22 | AFRICA |
| Pygmy_Bongo_GabS | 24 | AFRICA |
| Pygmy_Nzebi_Gab | 20 | AFRICA |
| Pygmy_Nzime_Cam | 52 | AFRICA |
| STU | 96 | 1000 GENOMES |
| TSI | 106 | 1000 GENOMES |
| Ust_Ishim | 1 | ANCIENT EUROPEAN |
| Wichi | 19 | SOUTH AMERICA |

| | | |
|---|---|---|
| YRI | 101 | 1000 GENOMES |

**Supplementary Table 2 – Summary genetic data for 20 EHPNG linguistic groups ordered identically to $F_{ST}$ table (Supplementary Figure 2)**

| GROUP | HETEROZYGOSITY | RUNS OF HOMOZYGOSITY | AVG LD | RELATEDNESS |
|---|---|---|---|---|
| ANGA | 0.205 | 3.375 | 0.167 | 0.039 |
| PAWAIAN | 0.205 | 3.846 | 0.164 | 0.073 |
| TAIRORA | 0.205 | 3.313 | 0.168 | 0.055 |
| AUYANA | 0.208 | 0.938 | 0.162 | 0.064 |
| AWA | 0.205 | 1.563 | 0.173 | 0.056 |
| GADSUP | 0.213 | 0.667 | 0.152 | 0.057 |
| AGARABI | 0.211 | 1.188 | 0.173 | 0.057 |
| KAMANO | 0.211 | 1.125 | 0.154 | 0.054 |
| SFORE | 0.214 | 1.125 | 0.152 | 0.044 |
| GIMI | 0.214 | 1.188 | 0.149 | 0.050 |
| NFORE | 0.214 | 0.438 | 0.155 | 0.058 |
| KEIAGANA | 0.214 | 0.938 | 0.148 | 0.055 |
| YATE | 0.209 | 2.313 | 0.144 | 0.049 |
| YAGARIA | 0.212 | 1.375 | 0.151 | 0.060 |
| BENABENA | 0.213 | 0.867 | 0.145 | 0.038 |
| LABOGAI | 0.216 | 0.313 | 0.145 | 0.057 |
| YABIYUFA | 0.215 | 0.250 | 0.140 | 0.053 |
| SIANE | 0.215 | 0.333 | 0.146 | 0.059 |
| GAHUKU | 0.215 | 0.333 | 0.145 | 0.052 |
| ASARO | 0.215 | 0.375 | 0.147 | 0.051 |

**Supplementary Table 3 – Breakdown of FS clustering in Village Analysis dataset; K=13**

| | FS CLUSTER | | | | | | | | | | | | |
|---|---|---|---|---|---|---|---|---|---|---|---|---|---|
| | **1** | **2** | **3** | **4** | **5** | **6** | **7** | **8** | **9** | **10** | **11** | **12** | **13** |
| **DESCRIPTOR** | **(Anga)** | **(Pawaian)** | **(Tairora)** | **(Eastern)** | **(SFore - Pamasugina)** | **(NFore)** | **(SFore - Atigina)** | **(Auyana)** | **(Awa)** | **(Northwest)** | **(BenaBena/Labogai)** | **(Gimi)** | **(Mid)** |
| **LINGUISTIC GROUP** | | | | | | | | | | | | | |
| ANGA | 22 | 0 | 0 | 0 | 0 | 0 | 0 | 0 | 0 | 0 | 0 | 0 | 0 |
| PAWAIAN | 0 | 13 | 0 | 0 | 2 | 1 | 0 | 0 | 0 | 0 | 0 | 0 | 0 |
| TAIRORA | 1 | 0 | 33 | 2 | 0 | 0 | 0 | 1 | 0 | 1 | 0 | 0 | 0 |
| AUYANA | 0 | 0 | 1 | 0 | 0 | 0 | 0 | 21 | 0 | 0 | 0 | 0 | 0 |
| AWA | 0 | 0 | 0 | 0 | 0 | 0 | 0 | 1 | 10 | 0 | 0 | 0 | 0 |
| GADSUP | 0 | 0 | 0 | 13 | 0 | 0 | 0 | 2 | 0 | 0 | 0 | 0 | 1 |
| AGARABI | 0 | 0 | 0 | 17 | 0 | 0 | 0 | 0 | 0 | 0 | 0 | 0 | 0 |
| KAMANO | 0 | 0 | 0 | 24 | 0 | 0 | 0 | 0 | 0 | 0 | 0 | 0 | 0 |
| SOUTHFORE | 2 | 1 | 0 | 4 | 177 | 17 | 101 | 3 | 20 | 7 | 3 | 9 | 19 |
| GIMI | 1 | 0 | 0 | 0 | 6 | 1 | 9 | 0 | 0 | 0 | 0 | 9 | 2 |
| NORTHFORE | 1 | 1 | 0 | 0 | 1 | 79 | 4 | 27 | 0 | 0 | 2 | 1 | 10 |
| KANITE | 0 | 0 | 0 | 1 | 0 | 0 | 0 | 0 | 0 | 0 | 0 | 0 | 2 |
| KEIAGANA | 0 | 0 | 0 | 0 | 0 | 2 | 0 | 0 | 0 | 0 | 0 | 0 | 25 |
| YATE | 0 | 0 | 0 | 1 | 0 | 0 | 0 | 1 | 0 | 0 | 0 | 0 | 18 |
| YAGARIA | 0 | 1 | 0 | 0 | 0 | 0 | 0 | 0 | 0 | 0 | 1 | 1 | 84 |
| BENABENA | 0 | 0 | 1 | 1 | 0 | 0 | 0 | 0 | 0 | 0 | 14 | 4 | 4 |
| LABOGAI | 0 | 0 | 0 | 0 | 0 | 0 | 0 | 0 | 0 | 7 | 12 | 0 | 0 |
| YABIYUFA | 0 | 0 | 1 | 0 | 0 | 0 | 0 | 0 | 0 | 18 | 3 | 0 | 0 |
| SIANE | 0 | 1 | 0 | 0 | 0 | 0 | 0 | 0 | 0 | 23 | 0 | 0 | 0 |
| GAHUKU | 0 | 1 | 0 | 0 | 0 | 0 | 0 | 0 | 0 | 11 | 4 | 0 | 0 |
| ASARO | 0 | 0 | 0 | 0 | 0 | 0 | 0 | 0 | 0 | 16 | 1 | 1 | 0 |

**Supplementary Table 4 – Summary of SOURCEFIND Results – All populations acting as sources (top), only external populations (bottom)**

| CLUSTER | LINGUISTIC GROUP COMPOSITION | CLUSTER_1 | CLUSTER_2 | CLUSTER_3 | CLUSTER_4 | CLUSTER_5 | CLUSTER_6 | CLUSTER_7 | CLUSTER_ | CLUSTER_9 | CLUSTER_10 | CLUSTER_11 | CLUSTER_12 | CLUSTER_13 | EXTERNAL_CLUSTER_23 | EXTERNAL_CLUSTER_25 | EXTERNAL_CLUSTER_53 |
|---|---|---|---|---|---|---|---|---|---|---|---|---|---|---|---|---|---|
| EHPNG CLUSTER 1 | ANGA | 0% | 0% | 0% | 0% | 0% | 0% | 0% | 0% | 10% | 0% | 0% | 0% | 0% | 0% | 90% | 0% |
| EHPNG CLUSTER 2 | PAWAIAN | 0% | 0% | 0% | 0% | 0% | 0% | 0% | 0% | 0% | 0% | 0% | 0% | 0% | 100% | 0% | 0% |
| EHPNG CLUSTER 3 | TAIRORA | 0% | 0% | 0% | 100% | 0% | 0% | 0% | 0% | 0% | 0% | 0% | 0% | 0% | 0% | 0% | 0% |
| EHPNG CLUSTER 4 | GADSUP, AGARABI, KAMANO | 0% | 0% | 49% | 0% | 0% | 0% | 0% | 21% | 0% | 0% | 11% | 0% | 16% | 0% | 3% | 0% |
| EHPNG CLUSTER 5 | SOUTH FORE - PAMASUGINA | 0% | 0% | 0% | 0% | 0% | 0% | 96% | 0% | 4% | 0% | 0% | 0% | 0% | 0% | 0% | 0% |
| EHPNG CLUSTER 6 | NORTH FORE | 0% | 0% | 0% | 0% | 0% | 0% | 31% | 32% | 0% | 0% | 0% | 0% | 36% | 0% | 0% | 0% |
| EHPNG CLUSTER 7 | SOUTH FORE - ATIGINA | 0% | 0% | 0% | 0% | 38% | 28% | 0% | 0% | 3% | 0% | 0% | 30% | 0% | 0% | 0% | 0% |
| EHPNG CLUSTER 8 | AUYANA | 0% | 0% | 6% | 14% | 0% | 49% | 0% | 0% | 31% | 0% | 0% | 0% | 0% | 0% | 0% | 0% |
| EHPNG CLUSTER 9 | AWA | 3% | 0% | 1% | 0% | 36% | 0% | 0% | 55% | 0% | 0% | 0% | 0% | 0% | 0% | 4% | 0% |
| EHPNG CLUSTER 10 | YABIYUFA, SIANE, GAHUKU, ASARO | 0% | 0% | 0% | 0% | 0% | 0% | 0% | 0% | 0% | 0% | 49% | 0% | 0% | 0% | 0% | 51% |
| EHPNG CLUSTER 11 | BENABENA, LABOGAI | 0% | 0% | 0% | 3% | 0% | 0% | 0% | 0% | 0% | 52% | 0% | 17% | 27% | 0% | 0% | 0% |
| EHPNG CLUSTER 12 | GIMI | 1% | 0% | 0% | 0% | 11% | 0% | 41% | 0% | 0% | 0% | 30% | 0% | 16% | 0% | 0% | 0% |
| EHPNG CLUSTER 13 | YAGARIA, YATE, KEIAGANA, KANITE | 0% | 0% | 0% | 2% | 0% | 30% | 0% | 0% | 0% | 0% | 56% | 11% | 0% | 0% | 0% | 0% |

| EHPNG target | Linguistic group composition | posterior.prob | CENTRAL_HULA_KEAPARA | CENTRAL_KOIARI | CENTRAL_MOTU | CENTRAL_WAIMA | CHIMBU | ENGA | GULF | GULF_AKOYE | MADANG_COASTAL | MADANG_HIGH | SHIGH_HULI | SHIGHLANDS | WHIGH_KIWAI |
|---|---|---|---|---|---|---|---|---|---|---|---|---|---|---|---|
| CLUSTER_1 | Anga | -14772.94826 | 0 | 0 | 0 | 0 | 0 | 0 | 0 | 1 | 0 | 0 | 0 | 0 | 0 |
| CLUSTER_2 | Pawaian | -22096.89418 | 0 | 0 | 0 | 0 | 0 | 0 | 0 | 0 | 0 | 0 | 0 | 0 | 1 |
| CLUSTER_3 | Tairora | -22657.77703 | 0 | 0 | 0 | 0 | 0 | 0 | 0 | 0 | 0 | 0 | 0 | 0 | 1 |
| CLUSTER_4 | Agarabi, Gadsup | -19892.04901 | 0 | 0 | 0 | 0 | 0 | 0 | 0 | 0 | 0 | 0 | 0 | 0 | 1 |
| CLUSTER_5 | South Fore - Pamasugina | -16147.90055 | 0 | 0 | 0 | 0 | 0 | 0 | 0 | 0 | 0 | 0 | 0 | 0 | 1 |
| CLUSTER_6 | North Fore | -18071.98913 | 0 | 0 | 0 | 0 | 0 | 0 | 0 | 0 | 0 | 0 | 0 | 0 | 1 |
| CLUSTER_7 | South Fore - Atigina | -17512.93049 | 0 | 0 | 0 | 0 | 0 | 0 | 0 | 0 | 0 | 0 | 0 | 0 | 1 |
| CLUSTER_8 | Auyana | -20009.87047 | 0 | 0 | 0 | 0 | 0 | 0 | 0 | 0 | 0 | 0 | 0 | 0 | 1 |
| CLUSTER_9 | Awa | -20202.53554 | 0 | 0 | 0 | 0 | 0 | 0 | 0 | 0.38 | 0 | 0 | 0 | 0 | 0.62 |
| CLUSTER_10 | Siane, Asaro, Yabiyufa, Gahuku | -15886.74269 | 0 | 0 | 0 | 0 | 0.99 | 0 | 0 | 0 | 0 | 0 | 0 | 0 | 0.01 |
| CLUSTER_11 | BenaBena, Labogai | -18004.58467 | 0 | 0 | 0 | 0 | 0.48 | 0 | 0 | 0 | 0 | 0 | 0 | 0 | 0.52 |
| CLUSTER_12 | Gimi | -18641.87326 | 0 | 0 | 0 | 0 | 0 | 0 | 0 | 0 | 0 | 0 | 0 | 0 | 1 |
| CLUSTER_13 | Yagaria, Keiagana, Yate | -17135.61826 | 0 | 0 | 0 | 0 | 0 | 0 | 0 | 0 | 0 | 0 | 0 | 0 | 1 |

**Supplementary Table 5 – Summary of GLOBETROTTER Results**

| | | all populations | | |
|---|---|---|---|---|
| target | desc | gen time | r2 | combo |
| 10 | NW | 12.9 | 0.8 | chimbu and 11 |
| 11 | BenaBena/Labo | 8.9 | 0.66 | 10 and 13 |
| 13 | Keia/Yag/Yate | 7.3 | 0.74 | 11 and 6 |
| 6 | NF | 1.2 | 0.9 | 13and7 |

**Supplementary Table 6 – Number of EHPNG Individuals Used in ADMIXTURE analysis**

| **EHPNG GROUP** | **N** |
|---|---|
| KAMANO | 25 |
| ASARO | 25 |
| ANGA | 25 |
| NORTH FORE | 100 |
| SOUTH FORE | 100 |
| TAIRORA | 40 |
| GADSUP | 17 |
| YABIYUFA | 21 |
| SIANE | 21 |
| GAHUKU | 17 |
| YATE | 21 |
| PAWAIAN | 18 |
| AWA | 17 |
| AUYANA | 23 |
| GIMI | 29 |
| LABOGAI | 24 |
| YAGARIA | 79 |
| KEIAGANA | 26 |
| AGARABI | 26 |
| BENABENA | 26 |
